# Automated Design of Robust Genetic Circuits: Structural Variants and Parameter Uncertainty

**DOI:** 10.1101/2021.08.13.456094

**Authors:** Tobias Schladt, Nicolai Engelmann, Erik Kubaczka, Christian Hochberger, Heinz Koeppl

## Abstract

Genetic design automation methods for combinational circuits often rely on standard algorithms from electronic design automation in their circuit synthesis and technology mapping. However, those algorithms are domain-specific and are hence often not directly suitable for the biological context. In this work we identify aspects of those algorithms that require domain-adaptation. We first demonstrate that enumerating structural variants for a given Boolean specification allows us to find better performing circuits and that stochastic gate assignment methods need to be properly adjusted in order to find the best assignment. Second, we present a general circuit scoring scheme that accounts for the limited accuracy of biological device models including the variability across cells and show that circuits selected according to this score exhibit higher robustness with respect to parametric variations. If gate characteristics in a library are just given in terms of intervals, we provide means to efficiently propagate signals through such a circuit and compute corresponding scores. We demonstrate the novel design approach using the Cello gate library and 33 logic functions that were synthesized and implemented *in vivo* recently (*1*). We show that an average 1.3-fold and a peak 6.5-fold performance increase can be achieved by simply considering structural variants and that an average 1.8-fold and a peak 30-fold gain in the novel robustness score can be obtained when selecting circuits according to it.

**Graphical TOC Entry:** 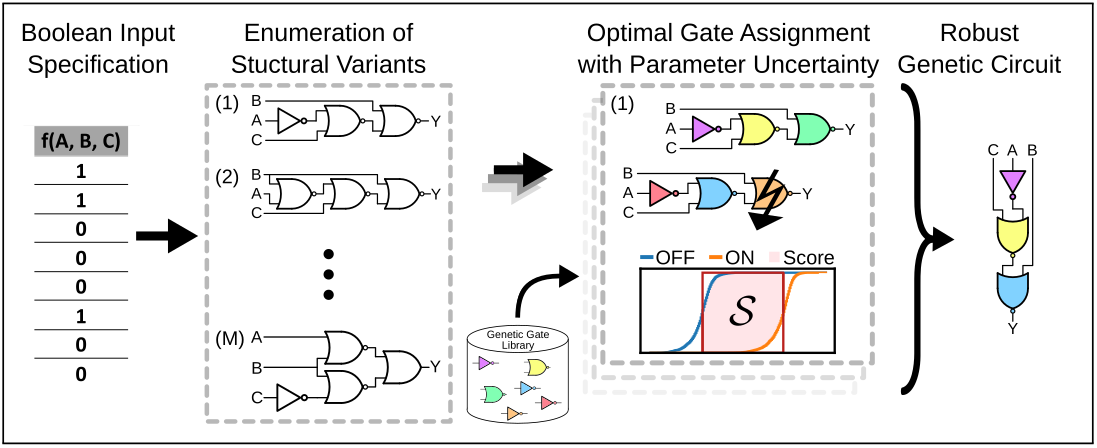

## 1 Introduction

Genetic design automation (GDA) parallels early efforts in electronic design automation (EDA) and recently also got to use state-of-the-art EDA tools to generate gene-regulatory circuits realizing combinational logic (*1*, *2*) as well as sequential logic (*3*). While historically EDA quickly ran into unmanageable computational complexity and hence devised clever approximate methods, current GDA problems are yet too small to require such approximations. In contrast to EDA’s scalability, GDA suffers from our limited understanding of what parameters fully characterize a genetic part or device (*4*–*6*) reflecting itself in GDA libraries with models of insufficient accuracy and scope. In particular, the context-dependency of circuit components (*7*) represents a central problem. That is, components behave differently depending on their adjacent up and downstream DNA sequences (*8*, *9*), on the specific resource allocation of the host organism (*10*, *11*), on the cross-talk from native regulatory factors (*12*, *13*) and on adjacent components that are biochemically up and downstream of the circuit (*14*, *15*). Cell-to-cell variability – referring to the fact that even within an isogenic cell population a synthetic circuit will behave differently from cell to cell – can also be understood as another context effect, i.e., the circuit functioning depends on the specific intracellular conditions realized within a particular cell. Cells may differ in their cell-cycle stage, their plasmid copy number and inevitably they will differ due to the random nature of biomolecular events, introducing copy number fluctuations in involved molecules (*16*, *17*). Such intrinsic noise will especially be important when the circuit is realized through lower abundant molecules, for instance through RNA regulators, (*18*, *19*), when compared to transcription factor based implementations.

As a consequence of cell-to-cell variability, the individual on and off expression levels for a genetic logic circuit may easily span one order of magnitude across a cell population (see e.g. (*1*)). For biomedical applications, such as disease detection and therapeutic circuits (*20*, *21*), stringent specifications are needed that guarantee the proper functioning of a circuit on the single-cell level and not just on bulk averages. As long as the on and off output levels cannot be assessed for each cell individually, such specifications translate to the requirement that the two distributions corresponding to the circuit’s on and off levels across the cell population, accessible for instance through flow-cytometry, do not show any overlap (*22*). In other applications such as biotechnology these requirements may be overly stringent and one is more concerned with just the fold-change between on and off bulk levels.

Taken together, current GDA tools such as Cello (*1*, *2*) require further domain specific adaptation in order to cope with context-dependency, the under-specification of part and device models and the intracellular variations encountered at the single-cell level. For instance, considering host energetics, GDA should find the circuit topology with the minimal number of components and should select the specific component realizations from the library that lead to robust circuits functioning under varying conditions. Existing tools for genetic circuit design (*23*) either use standard EDA methods and tools to determine the circuit topology, including Cello (*1*) and GeneTech (*24*), or leave the specification of the topology to the user and optimize inside its boundaries, like SBROME (*25*) does. iBioSim (*26*) uses an elaborate technology mapping algorithm that structurally matches library gates on a subject graph using branch-and-bound, but also constructs only one topology with minimal size in base pairs. Furthermore, Cello scores circuits based on the on and off levels corresponding to their median parametrization without incorporating variance information during the optimization process but provides predicting output distributions of the synthesized circuit. GeneTech doesn’t provide simulation capabilities, SBROME uses a deterministic gene expression model for single level output prediction only and iBioSim – while being very flexible in integrating simulation capabilities – couldn’t be found to incorporate simulation results in the synthesis and technology mapping process.

To this end, we propose the following extensions to the state-of-the-art GDA workflow. First, we demonstrate that better circuit topologies can be found compared to the ones obtained through generic EDA tools, exemplified by the 33 circuits reported in (*1*). We efficiently enumerate all structural circuit variants (*27*), which remains undoubtedly feasible for circuit sizes currently encountered in synthetic biology. Second, we improve the simulated annealing (SA) based gate assignment by employing neighborhood relation among all possible assignments (*28*–*30*). Since prominent placement tools for field programmable gate arrays (*31*) also utilize such neighborhood relation we adopted schemes from them. Third, we introduce parametric uncertainty in device models to mimic cell-to-cell variability, context-dependency or under-specification and extend the circuit scoring function to account for the incurred variability. We modify the traditional Wasserstein metric (*32*, *33*) to obtain a score that scales with the distance of the on and off levels and also reflects the degree of overlap among the corresponding distributions. Accordingly, two realizations of the same logic circuit showing same output medians across the complementary input assignments, and hence leading to identical scores in the traditional setting, could now be scored differently due to their possibly different output variability. Moreover, we develop a framework for robust design in the absence of probability distributions for specifying parametric uncertainty. In particular, if uncertainty is only given in terms of upper and lower bounds on the device parameters or gate characteristics we present a worst-case design approach based on envelope transfer function (see Fig. 1 for an overview).

**Figure 1:**
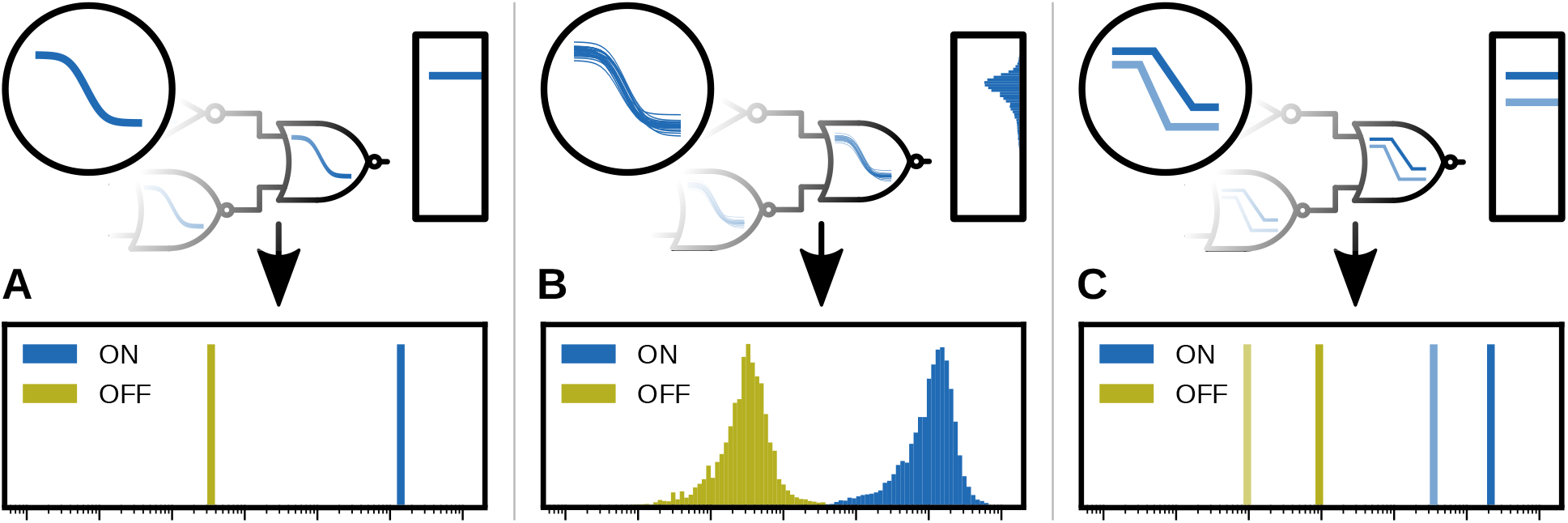
Different circuit design approaches. *A)* Traditional design and scoring approach with a nominal parametrization without uncertainty, as used by Cello (*1*). Cello does allow the prediction of output distributions but performs circuit synthesis only on median parametrizations; *B)* robust design approach accounting for cell-to-cell variability when probability distributions for device parameters are available, presented in this article; *C)* robust design solely based on interval specifications of transfer characteristics, presented in this article

## 2 Results and Discussion

### 2.1 General Problem Statement

This work deals with the particular problems of circuit synthesis and technology mapping in an automated generation of genetic logic circuits. It therefore focuses on jointly finding an optimal circuit topology *γ* in a set of topologies Γ and an optimal gate assignment *a* in a set of possible assignments 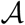 – which varies with the topology *γ* – given a library of gates 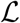 and a Boolean function specification 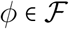. To formulate an optimization problem, we need a measure of compliance of a circuit (*γ, a*) with the functional requirement *ϕ*. This measure *S*(*γ, a*), which we call the circuit score, will be the optimization objective and we state the optimization problem as

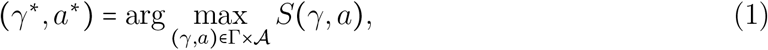

with the optimal topology *γ*^∗^ and assignment *a*^∗^. It is now crucial for the quality of the resulting logic circuit to take great care in specifying the set of possible topologies Γ on the one hand and the circuit score *S*(*γ, a*) on the other. In the following, we will discuss possible approaches to find and characterize application-optimal Γ and *S*(*γ, a*), which are compared with the approaches being part of the Cello framework (*1*). Since the dependence of 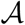 on the topology *γ* reflects the natural hierarchy of the problem, we will first address the synthesis problem and then proceed with the discussion on technology mapping and the score.

### 2.2 Circuit Synthesis involving Structural Variants

Prominent EDA tools, like ABC used in Cello, apply the cost functions area and delay (*34*), which are not directly suitable for genetic circuits, where fold-change and robustness pose the main challenges of design. We therefore enumerate circuits of all different topologies available from a given library of logic gates, which satisfy the logic function of the circuit. Since this structural enumeration is a combinatorial problem and quickly becomes infeasible, we optimize this procedure by following a hierarchical approach by considering only equivalent fan-out free circuits and performing pruning by isomorphism checking and the application of synthesis and library constraints online during enumeration (see Fig. 2A and also Method Section 4.2). After all fan-out free circuits have been found, we remove redundant gates inherent to this specific type of circuit topology to obtain the final set of circuits as generally structured Directed Acyclical Graphs (DAGs).

**Figure 2:**
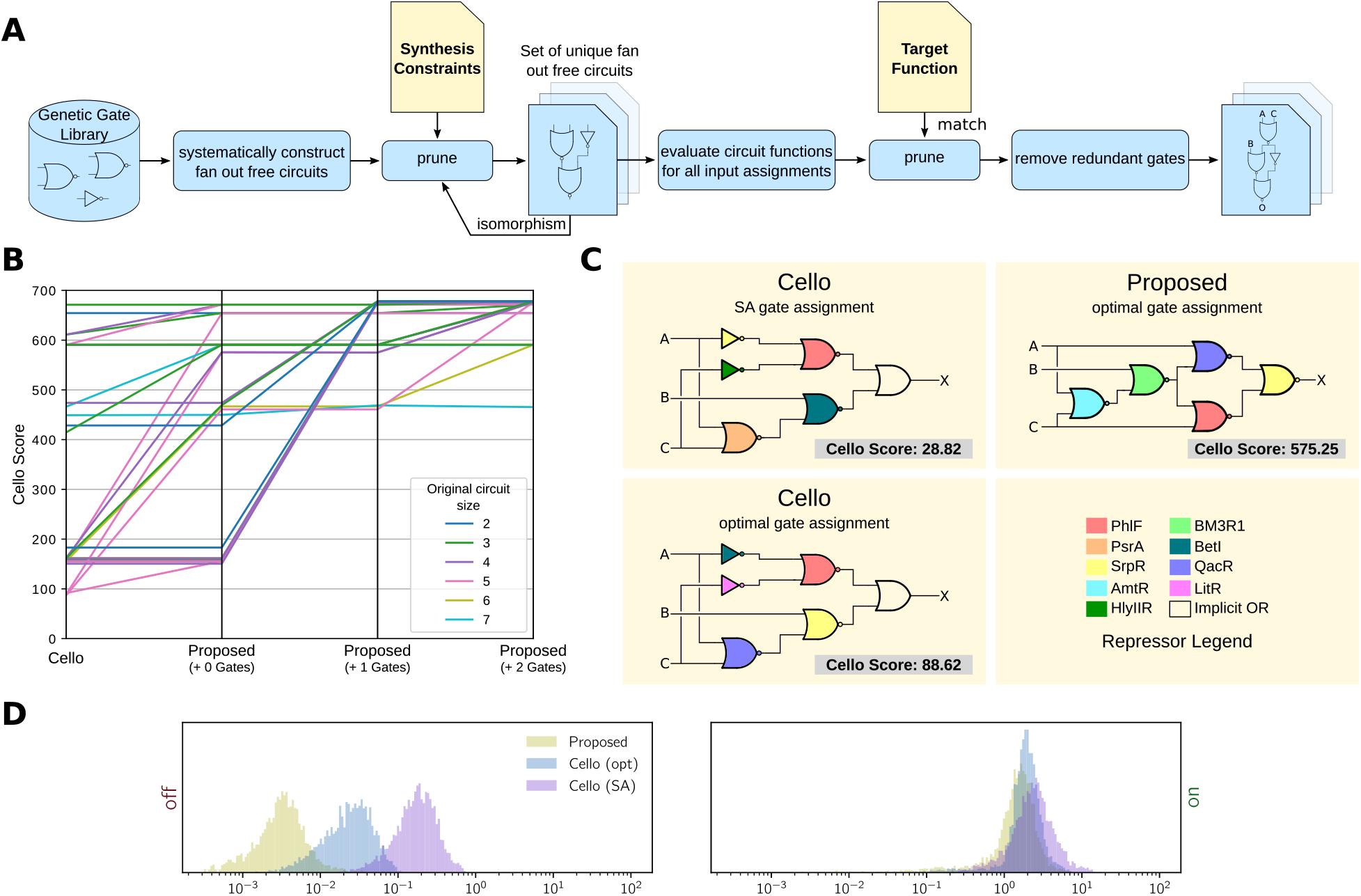
*A)* Synthesis flow for genetic circuits involving the enumeration of structural variants (also see Methods Section 4.2); *B)* Synthesis results for the 33 Boolean functions using Cello’s and our proposed synthesis approach with the number of excess gates allowed denoted in parentheses. Every function is represented by one line and its colour codes the size of its minimal circuit implementation. The monotonically ascending lines clearly show that the majority of circuits perform better using the proposed synthesis approach, while no circuit performs worse; *C)* Resulting circuits and their scores using Cello’s scoring metric for function 0x4D using Cello’s synthesis (with SA and optimal gate assignment) and our proposed synthesis approach. Given optimal gate assignments, the improved topology leads to a 6.5-fold improvement in the circuit score. Both circuit topologies feature the same number of genetic gates, as for the implicit output OR no physical realization is needed; *D)* Plot showing the output histograms of the circuits for function 0x4D. The proposed design features a lower output in the OFF case, thus increasing the separation between the complementary outputs and the Cello score.

In order to measure the benefit of including structural variety in genetic circuit synthesis, we synthesized all 33 functions provided in (*1*) using Cello’s library of genetic logic gates. In total, we carried out three runs of our proposed synthesis approach, constraining the search space differently. We only included circuits of minimum size in the first run and then relaxed this criterion to include one and two excess gates in the second and final run, respectively. At this point, we still used Cello’s circuit score metric to rate the separation of complementary Boolean outputs of the synthesized circuits. Finally, we compared our results to the circuits synthesized by Cello. To prevent fairness issues coming from Cello’s stochastic gate assignment optimization, we simulated all possible assignments exhaustively for both Cello’s and our circuit structures.

We found, that in the first run we were able to improve the circuit score of 14 of the examined 33 functions, while no circuit performed worse than the corresponding circuit synthesized by Cello and exactly the same number of logic gates was used (Fig. 2B). A 6.5-fold improvement in the score could be achieved maximally (Fig. 2C), while on average the scores improved by 28 %. Relaxing the considered circuit size to include up to one excess gate, the circuit score for 28 of the 33 functions could be improved up to 7.4-fold, leading to an overall improvement of 98 % on average compared to Cello. Relaxing the size by two excess gates, this trend continued (improvement for 31 of 33 functions up to 7.8-fold, 106 % on average). Thus, our synthesis approach not only improves on Cello for many of the considered functions using exactly the same number of logic gates, it also enables the designer to trade off circuit size against circuit performance deliberately (Fig. 2B). It also shows that genetic circuit synthesis profits from the additional degree of freedom of circuit topology. While the gate libraries are constricted and feature gates with heterogeneous transfer functions, it allows for placing well performing combinations of genetic gates in the circuit. For function 0x4D, for example, the proposed synthesis approach generated a circuit topology in which the output is driven by a NOR gate instead of the implicit OR gate while keeping the total number of genetic gates minimal (see. Fig. 2C). Fig. 2D depicts the increased separation of the complementary output states that leads to the improved Cello score of the proposed design.

### 2.3 Technology Mapping of Genetic Circuits using Neighborhood Heuristics

In EDA, the process of choosing logic gates from a library to implement a given circuit is called technology mapping (*35*). This process tries to find an assignment of gate realizations 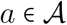 from the library 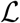 of real logic gates to the abstract logic gates in the circuit topology *γ* that optimizes a given score on the circuit. With regard to the presented circuit synthesis approach and the following statistical circuit evaluation method, an elaborate heuristic for technology mapping can contribute to alleviate the increased complexity in the synthesis process.

Cello already addresses the technology mapping problem with a generic Simulated Annealing (SA) heuristic to find the optimal gate assignment. However, since no problem specific knowledge is used during the generation of neighboring assignments by drawing gates from the library, their implementation can exhibit a far from optimal solution quality (see Fig. 2C). To alleviate this problem and obtain a more traversable assignment scoring landscape, we design a Markov policy for the random draws, which uses a metric that defines a distance between library gates on the space of analytical characteristics of the gates’ steady-state transfer functions (see Fig. 3A and also Method Section 4.3). Then a weighted euclidean distance in this space is used to allow drawing gates from an adaptive radius during SA (Fig. 3B, 3C).

**Figure 3:**
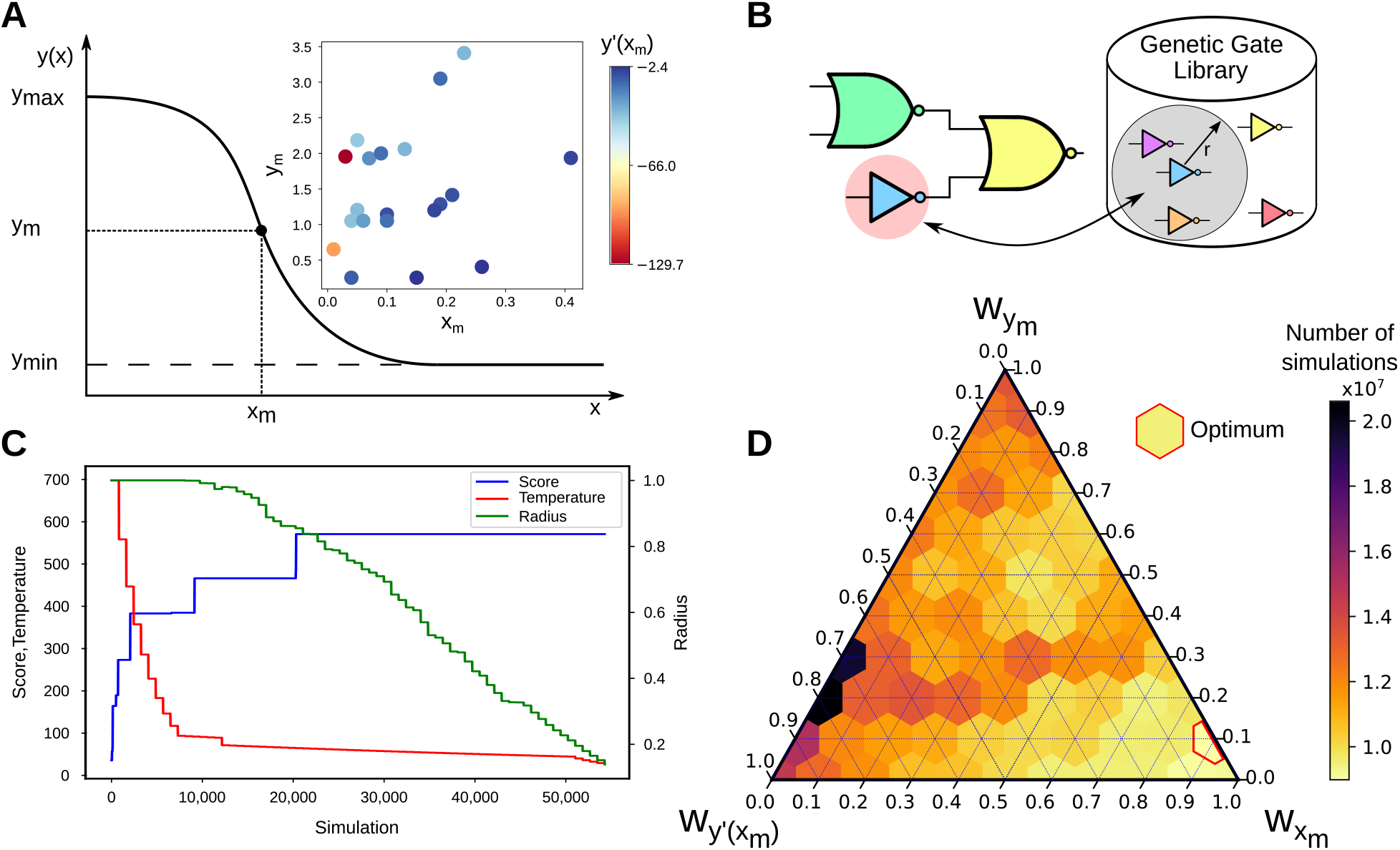
*A)* Parametrization of a general repressor Hill transfer function with offset and distribution of the considered genetic gates in the defined space of characteristics *x_m_*, *y_m_* and *y*′(*x_m_*); *B)* Radius based informed move of SA. The realization of one randomly selected gate of the circuit is swapped for a realization in the library based on the current radius *r*; *C)* Exemplary SA trace illustrating the adaptive radius; *D)* Number of simulations needed for mapping the set of benchmark circuits with SA applying 66 different weight configurations

To evaluate our technology mapping approach, we first compiled a set of 32 circuits by synthesizing multiple circuit variants for the Boolean functions examined in (*1*) and selecting circuits with 5 or more logic gates, thus sorting out circuits that are well assignable exhaustively. The problem sizes ranged from ~ 1 × 10^6^ to ~ 7.3 × 10^7^ possible gate assignments given the usage of Cello’s gate library. We then mapped the circuits using our basic SA and SA with proximity based neighborhood generation with different ratios of the distance weights. To account for SA’s stochastic run time, we repeated the mapping process 10 times and determined the mean run time of all runs.

Table 1 shows the mean score and number of simulations needed for different SA configurations compared to exhaustive search. Independently from the chosen weights, all SA runs yielded near-optimal scores. The base SA algorithm (no metric) reduced the number of simulations needed compared to exhaustive search by 97.5 %. Enabling the proximity based neighborhood generation with equally weighted dimensions, a further 1.61-fold speedup over basic SA is provided. For finding the best ratio of the weights given Cello’s gate library, we repeated the evaluation for the 66 different configurations depicted in Fig. 3D. Using the best configuration found, we were able to speed up the mapping process 2.23-fold across the set of 32 circuits and 5.8-fold for single circuits maximally over basic SA while still yielding near optimal technology mapping results. Mapping the benchmark set on a standard desktop PC, we measured a run time of 14.96 h for basic SA and 7.19 h using the best weight configuration.

**Table 1:** Mean number of simulations needed and mean score for different simulated annealing configurations across 32 circuits.

| Mapping Algorithm | Weight Config. | $w_{y_m}$ | $w_{x_m}$ | $w_{y'(x_m)}$ | Score | Simulations | Speedup |
| --- | --- | --- | --- | --- | --- | --- | --- |
| Exhaustive | – | – | – | – | 439.27 | 820,029,600 | 0.02 |
| SA | none | 0.0 | 0.0 | 0.0 | 439.18 | 20,475,365 | **1.0** |
| SA | equal | 1.0 | 1.0 | 1.0 | 439.00 | 12,696,430 | **1.61** |
| SA | best | 0.1 | 0.9 | 0.0 | 439.10 | 8,987,015 | **2.23** |

### 2.4 Robust Circuit Scoring

Signal propagation in genetic circuits varies significantly across members of a cell population due to context effects including those collectively termed cell-to-cell-variability. Therefore, a population-wide examination of such a circuit must naturally encompass a range of possible realizations of this circuit. We present two approaches to achieve such an inclusion. The first is based on a stochastic description of the circuit, which uses statistics of gate parametrizations and scores whole distributions of circuit outputs. The other is based on interval representations of transfer functions and signals to bound ranges of possible signal outputs of the circuit. Both approaches enter problem (1) by an appropriate choice of the score *S*(*γ, a*), which defines how we identify an optimal circuit and how much effort is needed to do so.

#### 2.4.1 Expectation-based Score (E-Score)

The score used by Cello is calculated using median realizations of the mapped gates’ known transfer function statistics, which are obtained empirically using flow cytometry measurements of isolated gates. Although this approach ignores the cell-to-cell variability of the circuit function, it results in a fast scoring procedure. While calculating any single circuit realization demands a similar runtime, the median realization is presumed to pose as what is deemed a typical realization of the respective circuit. However, this circumstance does not allow the user to trade computation time for scoring detail. To allow such a trade-off, we propose a sampling-based approach as an adjustable, parallelizable alternative, which – given an assignment – calculates output samples based on randomly drawn transfer function realizations from the known statistics and scores the resulting empirical distributions as a whole with a score, which roots in the Wasserstein distance (*32*). We can show, that the Wasserstein distance of the logarithmic output distributions emerges as a natural measure of separation corresponding to the population-wide expected on-off difference (see Methods 4.4.3). While the distance alone is a suitable candidate for comparing possibly overlapping output distributions in the sense of obtaining a functionally robust circuit, it is agnostic to variances in symmetric distributions. Although the obtained output distributions were often found to be skewed (in the direction of the complementary Boolean output), this insensitivity to variance is not suitable for a general score. We therefore chose to evaluate the distance partially in the sense depicted in Fig. 4B. We name the so obtained new score the E-score, and it allows us to score the negative impact of larger variance compared to an optimal output under a given median distance as shown in Fig. 4A and detailed in Methods 4.4.3. For the calculation in particular consider (8) in 4.4.3. Note, that as a consequence, the E-Score generally has a different absolute scale and a circuit scored by the E-Score is not necessarily comparable to one scored by the Cello score.

**Figure 4:**
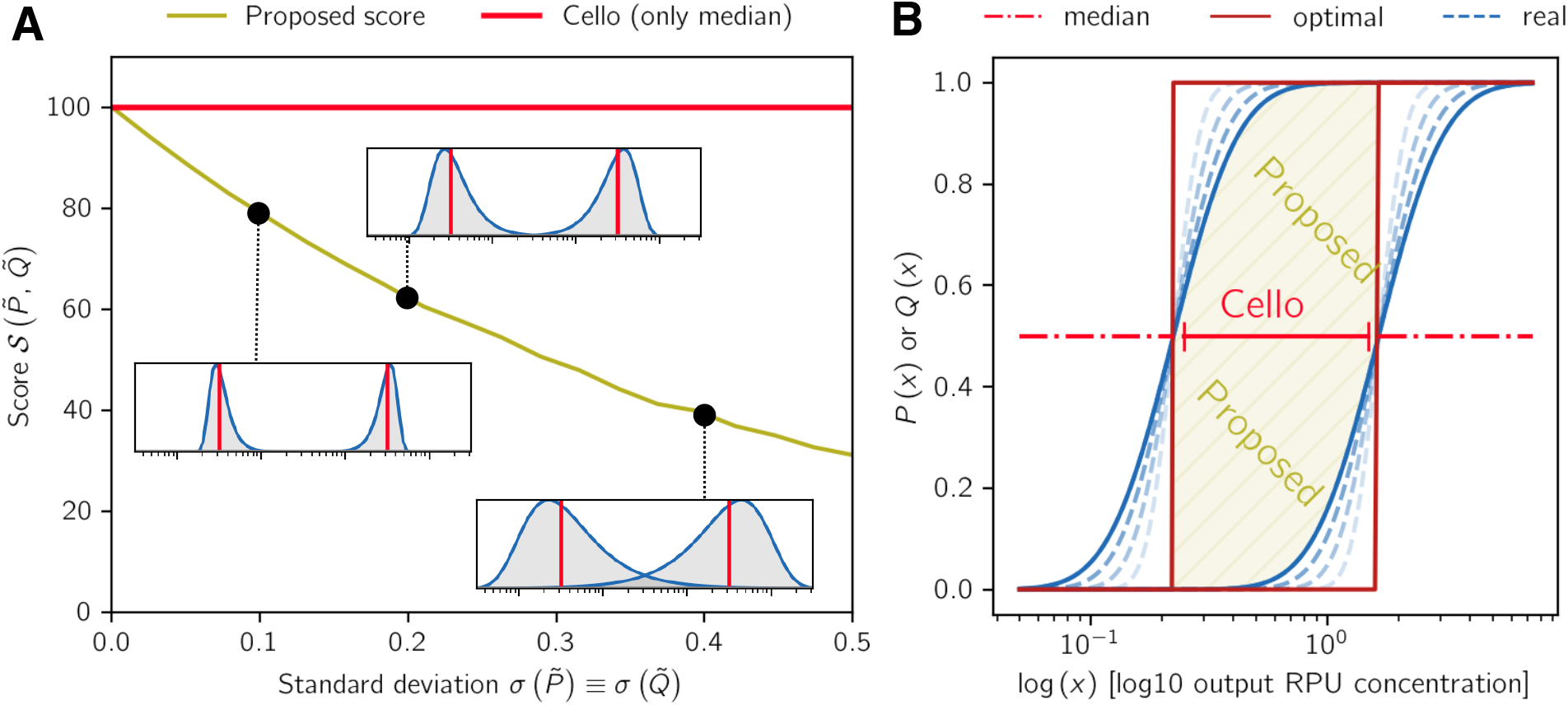
*A)* Proposed E-Score and Cello score of the two output distributions plotted over their standard deviation *σ*. The medians stay constant for all *σ*. Although intuitively the distributions with higher variance would be considered worse, Cello’s score doesn’t take this into account; *B)* Illustration of the two scores. The CDF’s of the two distributions representing Boolean on and off are plotted. An optimal output would concentrate all probability mass at specific points, which are considered to be at the median locations in accordance to Cello. Our score tries to capture the area enclosed by the inner tails of the output distributions within the optimal boundaries in the way hatched in gold, while Cello only builds the difference between two points. Choosing the Wasserstein-equivalent (cf. (7) in Methods 4.4.3) scores the area between the two blue lines, which would equal Cello’s score.

The sample realizations of the gate transfer functions themselves are obtained from sampled points of “noisy” Hill functions. These sampled points are obtained from Cello’s median realization processed together with histograms generated from flow cytometry data, which are sourced from Cello’s user constraint files (UCFs). Processing these has been done in accordance to the instructions from Cello’s supplementary material. The points are sampled, such that they represent equal quantiles on the so obtained empirical CDFs. We fitted Hill functions to these points, so that Cello’s median realization becomes a special case of a set of quantile realisations leading to empirical output distributions, which as a whole score the circuit (Fig. 4A and B). If we speak of quantile realizations, we mean these fitted gate transfer functions, which match specific quantiles on the empirical CDFs from Cello’s data. A more detailed description on how the samples have been obtained is given in Methods 4.4.2. To generate the circuit’s output distributions, first a sample circuit input is chosen. Then, an individual sample quantile realization is taken for each of the circuit’s gates and a circuit output sample is obtained from calculation of the circuit’s transfer function. This is done multiples times with new samples each time, until a desired refinement of the so obtained empirical output distribution is achieved. Details on the calculation are found in Methods 4.4.1

To test the procedure, we first rescored all circuits with ≤ 6 gates with their previous optimal assignments obtained from the exhaustive search using Cello’s original score described above, but this time drawing 5000 quantile realizations and using the E-Score. Unsurprisingly, since our score is stricter than the Cello score, the scores have been significantly lower (Fig. 4A). We kept the same circuit topologies obtained originally by Cello to retain comparability and only changed the gate assignment based on the new score. We found the best gate assignments for these topologies exhaustively while incorporating all sample realizations and the E-Score instead of only the median realizations and the original score. We could improve 21 of 31 assignments. The median improvement (only the improved assignments) was by 21.13% in score, while the mean improvement was at 179% (we will come back to this in a few sentences). If the circuit could be improved, on average 44.7% of the gates have been exchanged in comparison to Cello. The mean number of gates in improved circuits has been 5.39, while in kept circuits it has been 3.2. The reason for the large mean improvement is, that we could – using the histogram data – identify some possibly error prone circuits in Cello’s exhaustive results, which become erroneous under given variability. We use the term “erroneous” circuit here as a simplifying term for circuits, which result in a large fraction of inverted Boolean outputs using the sampled circuit realizations. We assume, the reason for such an erroneous behaviour can generally be found in subsequent alignments of the like depicted in Fig. 5. Since Cello’s score is agnostic to the distance of the median inputs to the transition regions of the gate’s transfer functions, a so chosen assignment might lead to inverted outputs in a real circuit where cell-to-cell variability is present. The E-Score aims to avoid such assignments. This lead in the extreme to a nearly 30-fold improvement in score in circuit 0x1C. The target output levels of 0x1C stayed unchanged, since the final gate has been kept. Additionally, to demonstrate the practicability of the SA heuristic in combination with the E-Score, we mapped the two largest circuits 0x41 and 0x81 with ~ 7.3 × 10^7^ possible assignments using SA and compared the results with the (exhaustively obtained) best possible assignments from Cello while still not modifying the circuit topology. Despite the stochastic optimization, both circuits could be improved (0x41 and 0x81 significantly by 84.9% and 40.92%). Exemplary output histograms for circuit 0x81 and the restored non-functional circuit 0x1C are given in Fig. 5C and D. We can conclude that, especially for high cell-to-cell variability, a higher confidence in the functionality of the so obtained circuit w.r.t a whole population can be achieved incorporating known statistics in the technology mapping process. To give an overview of the experiments, we provide statistical results in Table 2, where we compare sample scoring runs utilizing 5000, 500, 100, and 50 samples with the result obtained using Cello’s score. While excluding erroneous circuits (c.f. Fig. 5D), our score was able to reduce the variance of the logarithmic output distributions significantly as well.

**Figure 5:**
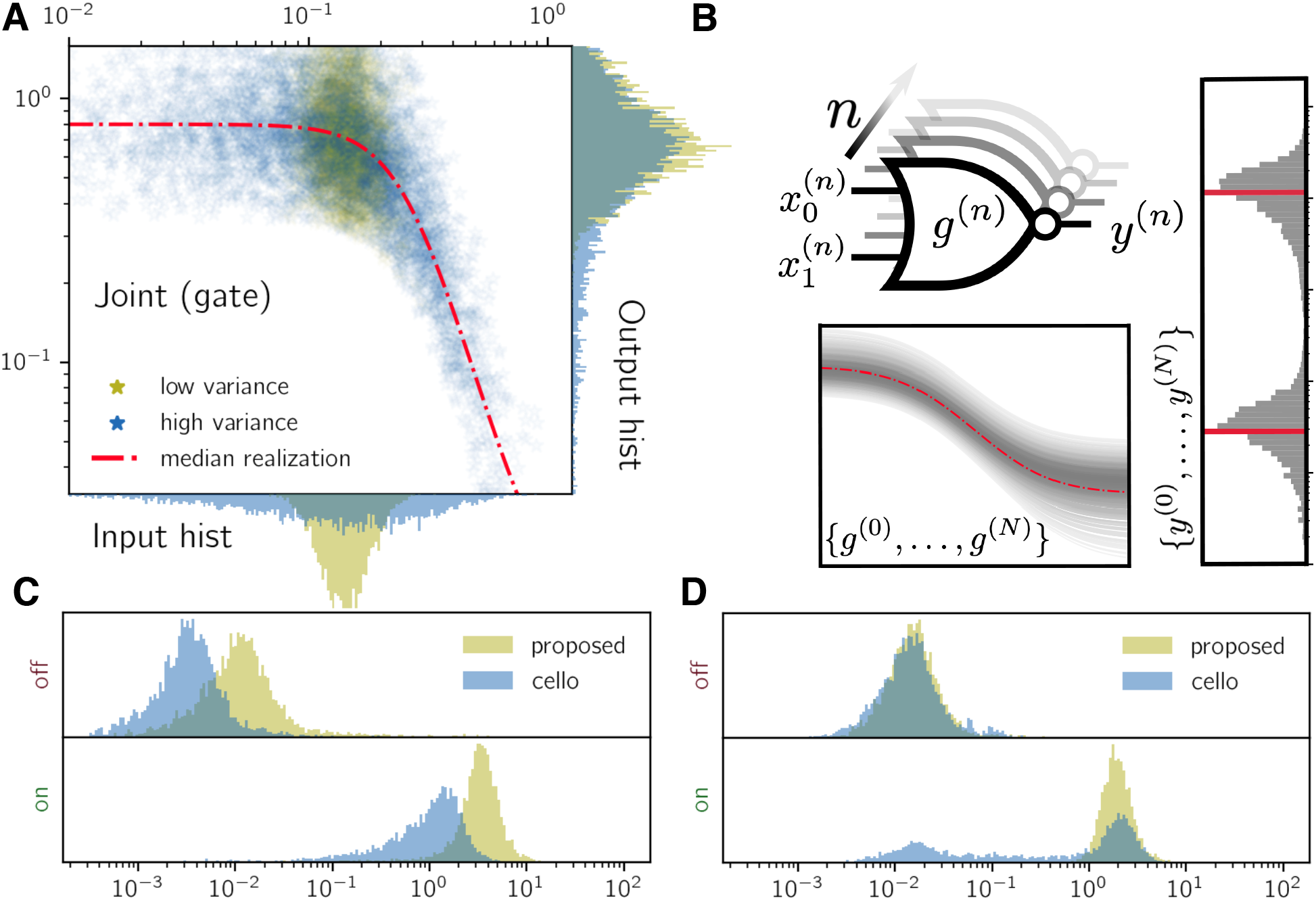
*A)* Input, output and joint histograms for a sample gate I/O scenario. The gate corresponds to promoter ‘BM3R1’ with ribosome binding site ‘B3’. The gates transfer statistics are reconstructed using the flow cytometry data from the Cello UCF ‘Eco1C1G1T1.pAN1201.UCF.json’. If only the medians of input distributions and gate transfer functions are considered like in Cello, the blue output would be considered a better result compared to the yellow one; *B)* Illustration of the sampling procedure. *N* parametrizations are pre-drawn for each gate for the respective environment and combined under independence assumption to yield the circuit output; *C) and D)* Plot showing the two histograms generated for the best assignments chosen by the respective scoring scheme. C: 0x81 and D: 0x1C. The optimal assignment of circuit 0x1C under Cello score results in many inverted Boolean outputs with given cell-to-cell variability and under the independence assumption made for the sampling.

**Table 2:**
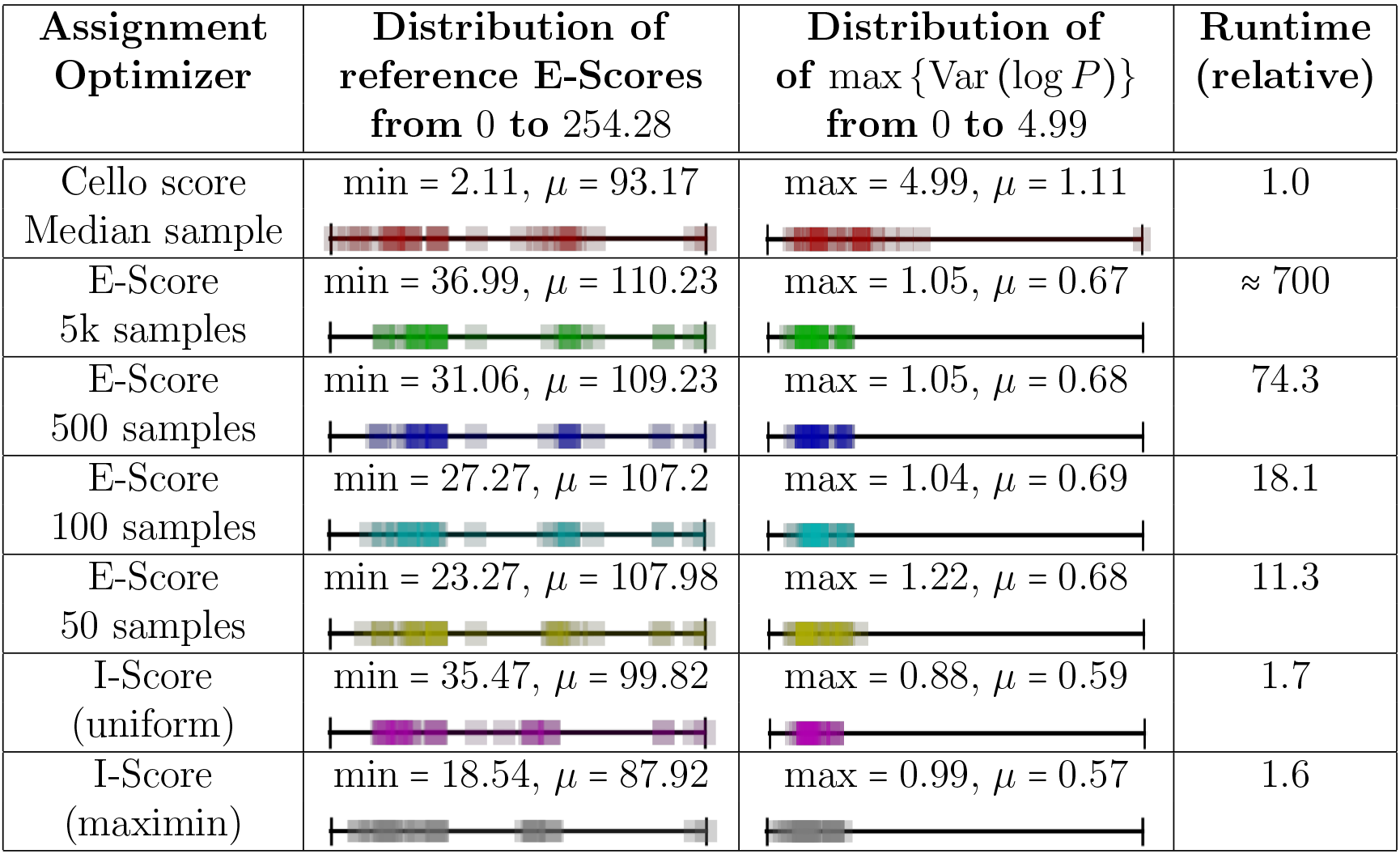
Table listing the exhaustive runs (31 circuits) giving an impression of different scoring schemes (Cello score, E-Score (5000 samples – used as a reference), E-Score (500 samples), E-Score (100 samples), E-Score (50 samples), I-Score (uniform), I-Score (maximin)). The median reference E-Score was roughly the same ≈ 73 for all. Besides the reference score with 5000 samples, which incorporates the most detail of the output distributions among all scores presented, we used the maximum variance of the logarithmic output distributions as another measure of fitness for the resulting assignment. We remember the calculations (8) for the E-Score and (11) for the I-Score

| Assignment Optimizer | Distribution of reference E-Scores from 0 to 254.28 | Distribution of $\max \{ \text{Var}(\log P) \}$ from 0 to 4.99 | Runtime (relative) |
| --- | --- | --- | --- |
| Cello score<br>Median sample | min = 2.11, $\mu = 93.17$ | max = 4.99, $\mu = 1.11$ | 1.0 |
| E-Score<br>5k samples | min = 36.99, $\mu = 110.23$ | max = 1.05, $\mu = 0.67$ | $\approx 700$ |
| E-Score<br>500 samples | min = 31.06, $\mu = 109.23$ | max = 1.05, $\mu = 0.68$ | 74.3 |
| E-Score<br>100 samples | min = 27.27, $\mu = 107.2$ | max = 1.04, $\mu = 0.69$ | 18.1 |
| E-Score<br>50 samples | min = 23.27, $\mu = 107.98$ | max = 1.22, $\mu = 0.68$ | 11.3 |
| I-Score<br>(uniform) | min = 35.47, $\mu = 99.82$ | max = 0.88, $\mu = 0.59$ | 1.7 |
| I-Score<br>(maximin) | min = 18.54, $\mu = 87.92$ | max = 0.99, $\mu = 0.57$ | 1.6 |

#### 2.4.2 Interval-based Score (I-Score)

The E-Score uses inverse transform sampling to draw samples representing random quantiles on the histograms obtained from flow cytometry. While for an acceptable amount of samples and under correct assumptions this approach is versatile and guaranteed to provide a consistent result, it might be useful to think about efficient alternatives with a stronger focus on robustness. We present two such efficient alternatives based on interval estimation. We call these variants I-Score. One of the two variants implements the maximin principle fundamental to robust optimization (*36*) the other is based on inscribed distributions. Though by construction not able to express output separation tendencies in proportions of the population, the score is able to identify assignments, which shift at least one individual to wrong outputs or in proximity to possible decision boundaries. Details can be found in Methods 4.5, but we give a short summary in the following. The basis of this score are bounding envelopes derived from our set of estimated context parameters, which enclose all or almost all of the known gate transfer function realizations. We then create a modified circuit double in size to the original, which is able to propagate (interval bounded) signals through the enveloped circuit and generate output intervals, which bound the output signals of the whole population, see Fig. 6A. Scoring by the maximin principle on these intervals is then performed by taking the distance of the smallest lower interval boundary corresponding to Boolean 1 and the largest upper boundary corresponding to Boolean 0 (c.f. (11) in Methods 4.5). An illustration of this idea is given in fig. 6. Having obtained the output intervals, scoring by the maximin approach is just one among a variety of possibilities. As an example, we could as well suspect these output intervals to support distributions of output values again like in section 2.4.1. By having no additional information, a maximum entropy assumption – and therefore uniform distributions on the support enclosed by the output intervals – would be a reasonable choice, which we briefly refer to by uniform I-Score.

**Figure 6:**
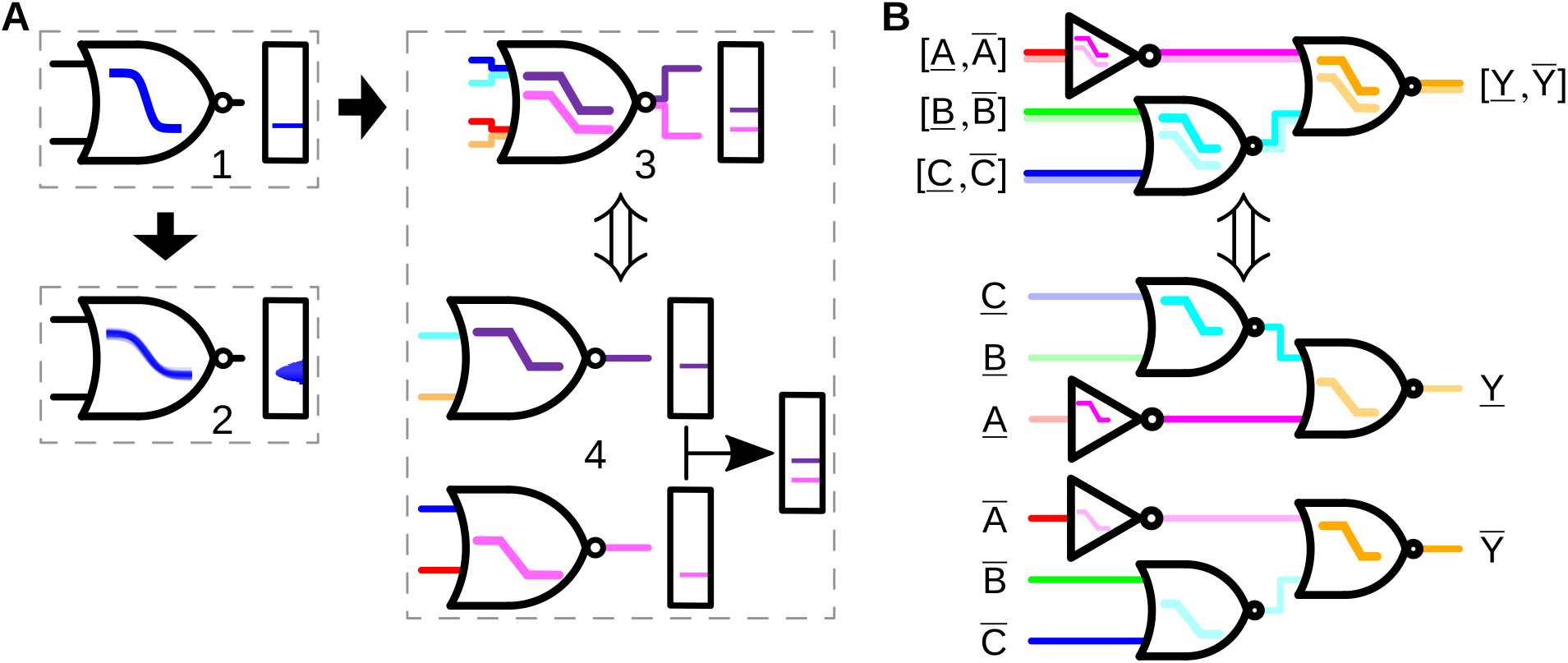
*A)* Overview of the designs considered within this work. The black arrows illustrate the direction of increasingly refined modelling; *A.1)* Cello scoring: model representing median parametrization without considering uncertainty; *A.2)* Expectation-based scoring (E-Score, eq. (8)): distributional information provided by parameter statistics taken into account; *A.3)* Interval-based scoring (I-Score, eq. (11)): enveloped model of transfer functions, consisting of a lower and upper envelope; *A.4)* Modified envelope-free circuit equivalent to the one shown in *A.3)*; *B)* Exemplary illustration of an enveloped circuit and its envelope-free version below. Note, that the wires in the enveloped circuit carry intervals and not scalar values, which is alleviated in the equivalent envelope-free circuit.

To evaluate the maximin approach, we again mapped all circuits with ≤ 6 gates using this score as a maximizer. We then rescored all circuits and their worst-case optimal assignments obtained with the maximin I-Score again using the expectation-based E-Score with 5000 quantile realizations. In comparison to Cello, of the 31 circuits 9 have been improved, 4 have been kept, and 18 have been worsened w.r.t the E-Score. The mean E-Score was the lowest of all tested scoring schemes, and as expected, the very bad E-Scores assumed by the Cello solutions have been avoided. Remarkable is the maximal variance of the logarithmic outputs. Their maximum has with 0.99 been significantly lower compared to Cello and also to some degree compared to the expectation-based scoring schemes. The mean maximal variance at 0.57 has been the lowest throughout. We then did the same experiment again with the only difference being, that we didn’t use the maximin I-Score on the output intervals but inscribed uniform distributions into these intervals and scored them using the E-Score. In comparison to Cello, of the 31 circuits 15 have been improved, 5 have been kept, and 11 have been worsened. The mean uniform I-Score has been around 4 points larger than that of Cello, while a very good minimum could be reached comparable to that of the full sampling E-Scoring. The maximal variance of the logarithmic outputs has been low overall as well. Its maximum has been the lowest throughout and its mean lies only a small portion above that of the stricter maximin approach.

Both schemes avoid erroneous circuits (large fraction of inverted Boolean outputs) and reduce output distribution overlap. Since the focus of the approach with inscribed uniform distributions on population-wide output separation is stronger, its minimal score has been almost as large as that of the baseline. Both interval-based approaches take less than two times the runtime of the Cello score, which has been the fastest overall. Unsurprisingly, the two interval-based scoring approaches also lead to output distributions with minimal log-variance. Like above, an overview can be found in table 2.

## 3 Conclusions

This work provides improvements to the emerging domain of genetic design automation, in particular for the synthesis of combinational logic circuits. We show that there is currently little need to make aggressive approximations in the circuit synthesis and the technology mapping step when compared to electronic design automation. Neither the implementable logic circuits nor the device libraries reach sizes that would require them. Using 33 example circuits from (*1*) we demonstrate that enumerating structural variants for a given Boolean specification and having an optimized stochastic search strategy in the technology mapping yield significantly better circuit realizations with an up to 27-fold improvement, all based on the traditional Cello library and scoring scheme (see Fig. 2). Under optimal gate assignments a 6.5-fold improvement can be achieved just due to structural variants, whereas for a given circuit structure one can find better gate assignments through a fast stochastic search that reliably finds the best assignment with a 2.2-fold speed-up (Table 1). Compared to the invested experimental time to actually implement and test genetic circuits, the incurred higher runtime for enumerating structural variants is negligible.

Going beyond those direct improvements of the established design process, the work presents a more general design approach that takes into account unavoidable underspecifications within biological device libraries, context-effects and cell-to-cell variability of circuit function. We show that accounting for them in the simplest way through parametric uncertainty, the design process yields more robust circuits, quantified in terms of a novel scoring metric that penalizes variance and overlap of the complementary circuit output distributions. We use random parametric families of Hill curves, learned directly from flow-cytometry data as gate models in the library and establish a fast Monte Carlo based scoring scheme. If uncertainty is only specified in terms of interval boundary, we provide another robust scoring scheme that just works with envelopes of gate characteristics and does not require any sampling step. The general methodology developed in this paper is not bound to a particular gate library. For libraries involving gates other than NOT and NOR gates, the neighbor-hood heuristic in the gate assignment can be adapted using correspondingly other features of the gate response curves. The proposed interval propagation method (Fig. 6) works for all monotone gate characteristics.

We see the work as a first step towards the use of more fine grained device models and the development of domain-adapted logic synthesis and technology mapping tools. There are several more extensions that we foresee in order for computer-based design methods to reach the necessary predictive power to be routinely used in the lab. Context-effects such as host energetics will require a more detailed biophysical model for how gate characteristics change under different conditions. Even if a random parametrization can account for that to a zeroths order, it will require the incorporation of a correlation structure among parameters that will be induced by cellular confounders like the cell’s energy state. Another aspect that also generates interdependence among gates is cross-talk due to, for instance, off-target binding of involved regulators or polymerase readthroughs for adjacent expression units. Such interdependency asks for enriched device models in libraries but will open up new interesting computational challenges for the circuit synthesis. Methods that account for intrinsic noise and for temporal aspects even for combinational logic (*37*), such as rise times or simple reversibility of circuit responses, are also yet to be developed. Integrating the temporal properties of genetic circuits that are central for designing sequential logic circuits (*3*) into a consistent robust design and scoring framework is another challenge ahead.

## 4 Methods

### 4.1 Robust Circuit Synthesis and Technology Mapping

In the following, we introduce the optimization problem formally in more detail compared to Section 2 and then dedicate separate sections to circuit synthesis and technology mapping/scoring. Let thus 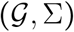 be the set of all labelled DAGs where 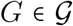 is a DAG with *G* = (*V, E*), *E* ⊆ *V* × *V* and labeling 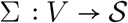 with 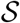 denoting the set of available types of functions (i.e. gate types) in that technology. Circuit synthesis returns a finite set of circuit topologies 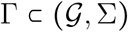 based on the synthesis map from the space of specifications in terms of Boolean formulae 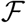 and an available library 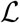, i.e., 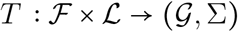. The technology mapping is the injective function *M* that takes each vertex of a topology *γ* in Γ and assigns it one element of library 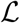, i.e., 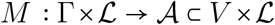. Both processes jointly result in a circuit (*γ, a*) with *γ* ∈ Γ and 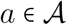. Rating such a circuit is then done using a circuit score function 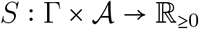 with the choice *S*(*γ, a*) exp (*S*(*γ, a*)), which we conveniently define to be the exponential of the log-score function 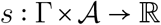. The definition of *S* as an exponential allows us to tackle the scoring in the logarithmic domain, which is more amenable with respect to the biological application. The score *S* is then quantifying the compliance of the circuit outputs with the Boolean functional requirement 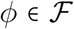. Proceeding from here, we can formulate the process of synthesis and technology mapping as an optimization problem of the form

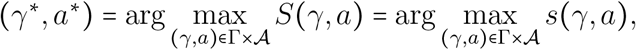

using the monotonicity of the logarithm for the last equality, with *γ*^∗^, *a*^∗^ being the optimal structure and assignment combination w.r.t the score *S*. The efficient construction of the set Γ and the proposed functional forms of *s* will be detailed in the following sections.

### 4.2 Circuit Synthesis involving Structural Variants

The problem of finding all structurally different implementations of a Boolean function is a DAG-enumeration problem. Thus, we intermediately enumerate all fan-out free circuit structures *C* = {*γ* ϵ Γ: ∀*v* ϵ *V*: |{*u* ϵ *V*: (*v, u*) ϵ *E*}| = 1}, simplifying enumeration and pruning (see Fig. 2A). During the systematic construction of *C* from the given set of gate types *S* in a library of genetic logic gates 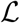 the found topologies are pruned according to the optional synthesis constraints maximum circuit weight *ω* and depth *δ*, i.e. ∀*γ* ϵ *C* ∶ |*γ*| ≤ *ω* ∧ *l* ≤ *δ*, with *l* being the longest path of *γ*. Furthermore, let *ϕ* be the *n*-ary Boolean target function and *I_γ_* = {*i*_0_, *i*_1_ …} be the set of unconnected gate inputs of *γ* then ∀*γ* ϵ *C*: |*I*_γ_ ≥ *n*. If the enumeration leads to isomorphism between the newly found topology *γ*′ and any existing topology *γ*, i.e. ∃*γ* ϵ *C*: *γ* ≃ *γ*′, *γ*′ is also discarded. The intermediate result is the complete set of unique fan-out free circuits consisting of gates of types *S* with a sufficient number of unconnected gate inputs to implement *ϕ*.

Then, a set of primary inputs 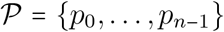 with 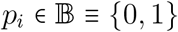 is instantiated and all possible assignments of unconnected gate inputs and primary inputs are generated, i.e. 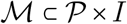. For each fully specified circuit the Boolean function is evaluated and thus the set of circuits *C_ϕ_* implementing *ϕ* is obtained, i.e. 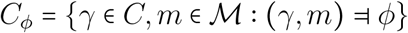. Redundant logic gates inherent to fan-out free circuits are then eliminated by evaluating their function w.r.t to the primary inputs and merging functionally equivalent gates, thus returning to a general DAG structure. This allows an application of final library constraints, i.e. checking whether the total number of genetic realizations in 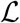 and the number of realizations per gate type 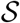 is sufficient to implement each circuit.

### 4.3 Technology Mapping of Genetic Circuits Using Neighborhood Heuristics

The smallest possible change that can be performed to generate a neighbor from a given solution is the substitution of one gate realization by another realization of the same logic type. Given that the gates, e.g., used in Cello differ greatly in their signal transfer behavior, a random substitution of one gate leads to an arbitrarily big change in the gate’s transfer function and thus in the circuit’s performance. Thus, we determine characteristic features of the gate realizations’ transfer functions and combine them into a proximity measure, enabling heuristic search algorithms to deliberately control the severity of changes to a solution during neighborhood generation.

The elementary transfer behavior of Cello’s genetic logic gates is characterized by a Hill repressor function

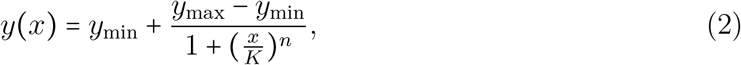

where *x* and *y* denote the input and output promoter activity, *y*_min_ and *y*_max_ define the output interval, *K* is the repression coefficient and *n* the Hill coefficient. This transfer function gives the gates a NOT or a NOR characteristic, depending on how many signals it is sensitive to. A feature used for characterizing electronic NOT gates is the switching threshold *V_m_*. It is defined as the point on the transfer function where *V*_in_ = *V*_out_ and impacts the device’s noise margins (*38*). Because of the global voltage levels *V*_DD_ and *V*_GND_ used commonly for input and output signals and thus symmetrical input and output intervals, *V_m_* can be found near the inverter curves inflection point for well built devices. Genetic logic gates lack a common reference value for input and output levels. Thus, we redefined the switching threshold for the considered genetic gates to be the point on the Hill curve, where an output concentration halfway between the minimum and maximum output concentrations is reached (see Fig. 3A). Let *y_m_* be that output concentration and *x_m_* the corresponding input concentration. We choose these characteristic features to be the first two dimensions of our proximity measure, i.e.,

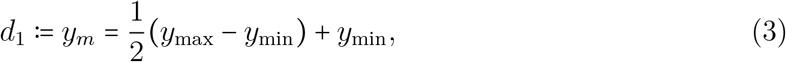

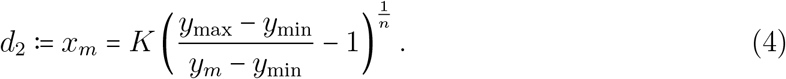

Further examination of the given gate library showed that the gates transfer functions differ greatly in the gradient at *y*(*x_m_*). Thus, we define the gradient *y*′(*x_m_*) at the switching threshold to be another characteristic feature

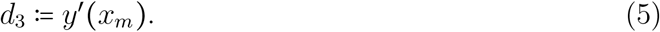

Denote by **d**_*i*_ the three-dimensional feature vector of gate *i* and define the diagonal weighting matrix **W** ϵ ℝ^3×3^ with entries *W_nm_* = *w_n_*/*δ_n_* for *n* = *m*, where *w_n_* ϵ [0, 1] is the adjustable weight for feature *n* (see Fig. 3D) and *δ_n_* the maximal absolute difference in the *n*-th feature between two gates across the whole library, then we can quantify the similarity between any two gates *i* and *j* in library by the **W**-norm

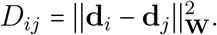

In order to evaluate if local search heuristics for the technology mapping of genetic circuits can benefit from the proposed proximity measure, we integrated it into the neighborhood generation of SA, that has been shown to profit from a well structured, problem specific neighborhood (*28*–*30*).

A major challenge when implementing SA is to specify central parameters like initial temperature and annealing schedule that lead to the desired solution quality and a reasonable run time. For the base implementation of the algorithm, we adopted these specifications from VPR, a tool for FPGA logic synthesis that uses SA for FPGA placement (*31*). Then, we adapted the algorithm to yield near-optimal results for the given technology mapping problem by slowing down the annealing schedule and conditioning the number of iterations per temperature level on the problem size. Here, the problem size is the number of possible gate assignments resulting combinatorially from the composition of gates in the circuit and in the library.

For every iteration *k*, VPR determines a radius *r_k_* in which logic cells on the chip are considered to be swapped in the search process. The ratio of the number of accepted solutions to the number of total evaluations *α* is calculated continuously during the annealing process and *r* is controlled to keep *α* near the empirically determined sweet spot of 0.44, i.e., *r_k_* = *r*_*k*−1_(1 − 0.44 + *α*). When, caused by the decreasing temperature, *α* drops below 0.44, the search radius *r* is decreased. This leads to a more local search for neighboring solutions in the late phase of the annealing process that are likely to have similar score values, thus leading to an increase of *α*. This ultimately results in the evaluation of less solutions with low scores that would be rejected anyway. We adapted this approach to our proximity based neighborhood generation. In our case, the radius controls which two gate realizations *i* and *j* in the library are considered for a swap, based on their distance *D_ij_*. The radius is initialized with the maximum distance of gates in the library, thus allowing for a global search in the search space in the early, high temperature phase. During the annealing process, *r* is decreased, progressively excluding gates with strongly differing transfer characteristics from the neighborhood generation. Further implementation details can be learned from the code available in a public repository.

### 4.4 Expectation-based Score (E-Score)

Like mentioned in section 2.4.1, to better represent the variability of the gates over different cellular contexts, considering statistical descriptions of the circuits and their outputs is one possible way. This improves the representation of population-wide circuit behaviour in the score function *S*(*γ, a*) (and therefore *S*(*γ, a*), which is used as a proxy). However, before we focus on the scoring in detail, we need a stochastic description of a genetic circuit. Therefore, we first introduce such a description, then we talk about how to generate sample realizations from this circuit, and finally, we talk about the score.

#### 4.4.1 Circuit Description respecting Cell-to-Cell Variability

Let thus 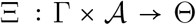 denote the parametrization of a circuit (*γ, a*). To represent the cellular context in terms of known statistics, we understand Ξ(*γ, a*) as a random variable characterized by a distribution Ξ(*γ, a*) ~ *P* (***θ***) associated with circuit (*γ, a*). In the following, if we speak of a circuit parametrization, a circuit realization or a specific context, we mean a particular realization Ξ(*γ, a*) = ***θ***, which we assume to be constant for each member in a population. Our goal will be to not only calculate the circuit output based on the median realization of the parameters, like Cello, but a set of sample outputs consistent with realizations based on the measured data, which jointly represent output distributions associated with a whole cell population. Since the circuit function under a fixed parametrization is – at this scale – assumed to be sufficiently deterministic, the output distributions depend on a vector of realizations representing the *M* circuit inputs 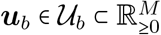 and the vector of realizations ***θ*** ϵ **Θ** representing the (cellular) context. Let further the realization of the random variable representing the 1-bit output be denoted by *v*. A Boolean label 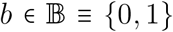 is attached to each set of input configurations 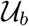 and its elements ***u***_*b*_ to indicate, which output *v* is associated with a Boolean value 0 or 1 from the truth-table *ϕ*. If we just write ***u***, we usually mean an arbitrary input without caring about any underlying logic function. The output density *p* (*v*) can be found by marginalization

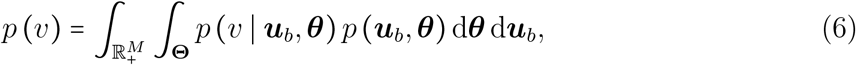

with *p*(*v* | ***u***_*b*_, ***θ***) being the density of the circuit output conditioned on a particular input and context realization. Given a gate library 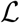 containing *L* context-dependent gate quasi-steady state transfer functions {*g*_1_, …, *g_L_*}, of which all are of a type 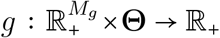, where *M_g_* is the number of gate inputs. Then, the circuit output can be calculated from a circuit transfer function *f*(***u***_*b*_, ***θ***) ≡ *f*(***u***_*b*_, ***θ***, *g*′, *g*′′, …) ≡ *f*(***u***_*b*_, ***θ***, *γ, a*) depending on the set of gates in the circuit 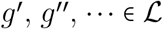. This circuit transfer function can be evaluated from subsequently calculating gate outputs. Therefore, the output conditional *p*(*v* | ***u***, ***θ***) can be calculated directly from *f*, since for a specific context ***θ*** and input realization ***u*** the circuits transfer function *f* is deterministic (as are all gates *g*). Consequently, *p*(*v* | ***u***, ***θ***) = *δ* (*v* − *f* (***u***, ***θ***)) is given by a degenerate distribution, where *δ* is the Dirac delta function. As a simplifying assumption, we require the factorizations *p*(***u***, ***θ***) ≡ *p*(***u***) *p*(***θ***) and 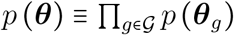. The first assumes input distributions independent of the cellular context and circuit chosen and the second, that the cellular context is acting independently on the gates in the circuit. This allows us to equip every gate with an individual set of sample realizations independent of which other gates are in the circuit. The latter enables initial sample generation for all gates in the library to allow a fast simulation in a technology mapping process. We require further, that *g_l_*(***x***, ***θ***) ≡ *g_l_*(***x***, ***θ***_*l*_) for all 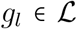 to allow learning the gate parameters from Cello’s isolated gate measurements.

Cello’s gate library has some properties, we need to address briefly. It consists only of NOT and NOR gates, where the latter combine multiple inputs to a single input via implicit summation. This means, if we write *g*(*x*, ***θ***), this also includes gates with *M_g_* > 1 by 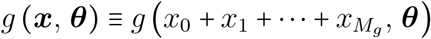, c.f. (*1*).

#### 4.4.2 Collecting Samples

We built our set of samples by taking the cytometry data from Cello’s UCFs. For each binned dataset in the UCF file associated with an input concentration from the discrete set *x* ≡ (*x*_0_, *x*_1_, …, *x_K_*) we define the empirical distribution 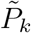 represented by the random variable 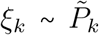, so that 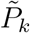 is represented by the binned dataset with its median logarithmically shifted to 0 (if not already). We multiplied these *ξ_k_* with the Hill functions representing median realizations 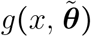 also present in the UCF file to obtain “noisy” hill function values 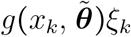 for each *k* (we added log(*ξ_k_*) in the logarithmic domain). We did this in accordance to the instructions from the Cello supplementary material. We thus obtain a new distribution 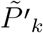 for each *k* with support logarithmically shifted by the constant 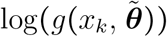. Employing inverse transform sampling, we drew a set of *N* iid standard uniform random variates ***q*** = (*q*_0_, *q*_1_, …, *q_N_*) representing quantiles and – using these and the inverses of the empirical CDFs – obtained *N* sets of *K* samples 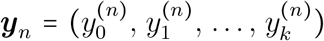 from the 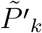 representing similar quantile locations for all the *k*. The relation between *q_n_* and 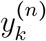 is then given by 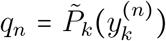. Let ***g***(***θ***) ≡ (*g* (*x*_0_, ***θ***), …, *g* (*x_K_*, ***θ***)) be the vector of gate outputs for each of the *x_k_* under realization ***θ***. We then solved the Tikhonov-regularized least squares regression problems 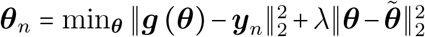 to obtain *N* sets of environment parameter samples ***θ***_*n*_ (we use Hill function parameters as a proxy) representing the variability captured by the cytometry measurements. Under the independence assumptions outlined in the previous section 4.4.1, we can generate the samples offline and store them in an extended gate library.

#### 4.4.3 The Score

Equipped with our definitions from above, we are now able to specify a suitable *S*(*γ, a*), which we use to score a context-dependent circuit. Like Cello, we use the logarithmic on-off difference as a basis for our score, which seems to be a suitable quantification of the separation of two values in the positive reals. However, in contrast to Cello, which calculates 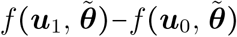 with the median realization 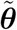, we have probability distributions to score if Ξ(*γ, a*) is a random variable. As a consequence *f* (***u***_1_, Ξ (*γ, a*)) − *f*(***u***_0_, Ξ (*γ, a*)) is a random variable as well. Therefore, we first chose its expectation as a scoring candidate, which manifests in the log-score

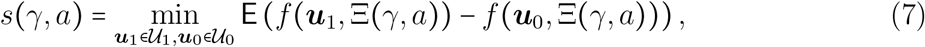

where 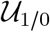 is the set of all real valued circuit input vectors associated with Boolean output 1/0 from the circuit’s truth-table *ϕ*. Let *f* (***u***_0_, Ξ (*γ, a*)) ~ *P*_0_ and *f* (***u***_1_, Ξ (*γ, a*)) ~ *P*_1_ for a specific ***u***_0_, ***u***_1_. So, *P*_0_ and *P*_1_ are the CDF’s of population-wide individual outputs associated Boolean 1 and 0 for specific circuit inputs ***u***_0_ and ***u***_1_. Then, interestingly, the expectation in (7) is equal to the Wasserstein distance of *P*_0_ and *P*_1_ if *P*_0_(*v*) − *P*_1_(*v*) never changes sign. This means, that looking at any arbitrary circuit output *v*′, there must lie more probability mass below this value associated with Boolean 0 than with Boolean 1, so *P*_0_(*v*′) > *P*_1_(*v*′). The Wasserstein distance, which is meant here, is defined on the metric space (ℝ_≥0_, |*x*_1_ − *x*_0_|) by

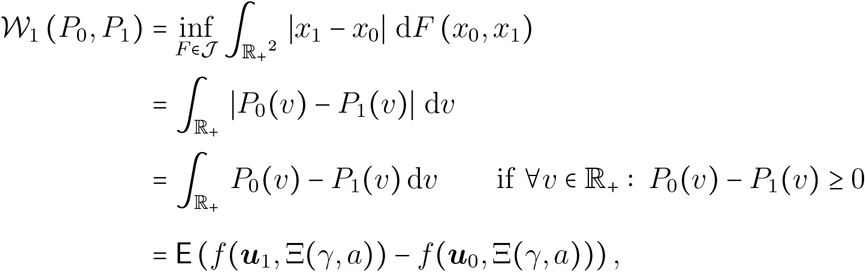

where 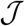 is the set of all joint probability measures *F* on 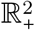, which have marginals *P*_0_ and *P*_1_. Note, that the last equality holds unconditionally. In our case, where we have two empirical distributions 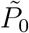 with samples 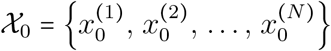 and 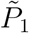 with samples 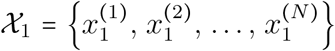, the calculation of *s*(*γ, a*) reduces to (cf. the analogy for 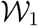 in (33))

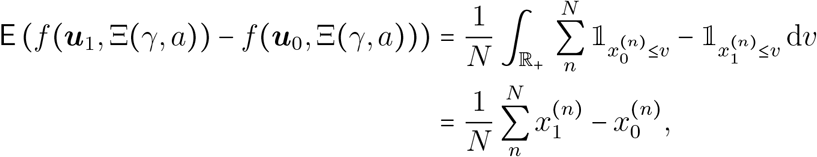

where 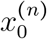 is the *n*-th order statistic (*n*-th smallest sample) in 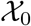. The same holds for 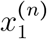 and 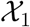. We discussed in Section 2.4.1 that, however, this score is agnostic to variance in symmetric distributions. Therefore, if the output distributions are symmetric, an overlap could not be detected. We therefore modify the score in the sense depicted in Fig. 4B to only score the negative deviation from a per-median optimal output window caused by the distributions’ variances. This formalizes in the log-score

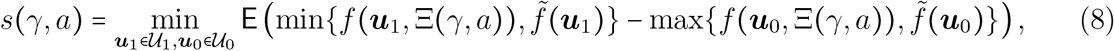

where 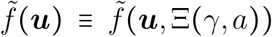 is the median circuit output for input ***u*** over Ξ(*γ, a*). We call the exponential *S*(*γ, a*) exp (*S*(*γ, a*)) with *S*(*γ, a*) from (8) the E-Score. Note, that this modification doesn’t reduce the computational effort in comparison to 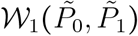 but doesn’t increase it notably either. The expectation in the score (8) can be calculated on the empirical output distributions by

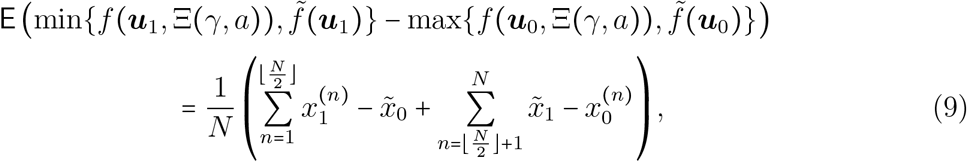

where 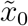 and 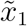 are the medians of 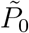 and 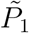. Note, that these are not equal to 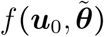 or 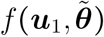, since the output of the median circuit realization does not guarantee to yield the median circuit output. Note, that the resulting score *S*(*γ, a*) generalizes Cello’s score. For degenerate distributions (two “samples”), it is simply given by *S*(*γ, a*) = exp (*x*_1_ − *x*_0_). In the case of Cello, the *x*_0_, *x*_1_ are the logarithms of the circuit outputs produced by the median realization 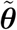 for two corresponding inputs ***u***_0_ and ***u***_1_.

### 4.5 Interval-based Score (I-Score)

Like mentioned in 2.4.2, we propose another approach, which is stricter and concentrates more on robust optimization (*39*). It is a consequent implementation of Wald’s maximin principle in the sense, that it doesn’t seek to negotiate the diversity of a population, like an expectation does, but find just the weakest element. This can also be the case, if we do not want to calculate samples to approximate an output distribution or do not have sufficient data to derive distributions of parameters. In this case, the circuit parametrization Ξ(*γ, a*) is not understood to be random anymore, but becomes a set-valued map, returning a set containing all known parameter realizations ***θ*** ϵ Ξ(*γ, a*) ⊂ **Θ** in circuit (*γ, a*). The associated maximin-score is then

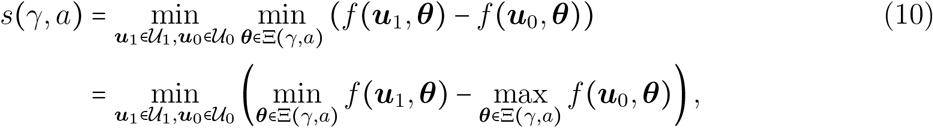

with an additional minimizer over the range of possible parameters. We now, without knowledge of existence, choose two parameter sets 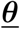 and 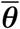, for which we demand the conditions, that for any 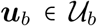 with 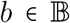 we have 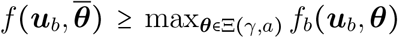 and 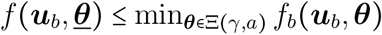 so that we obtain the following lower bound 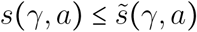

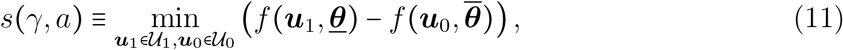

which we use as an interval-based score and call its exponential *S*(*γ, a*) = exp(*s*(*γ, a*) the I-Score. We can show, that if all gates in the circuit (*γ, a*) have transfer functions 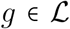 that are monotonous (either decreasing or increasing) for any fixed parametrization ***θ*** and 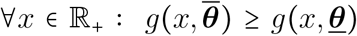, then 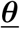 and 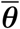 exist and the output intervals 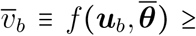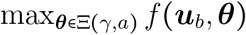 and 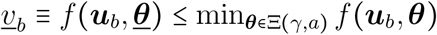 for 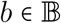 can be calculated only from the bounds 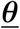 and 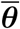. Since, like explained in 4.4.1, we use Cello’s gate library, which consists only of NOT gates and NOR gates with implicit summation, the monotonicity condition for all *g* is satisfied. Additionally, because we derived all available samples from Cello’s cytometry data and the bounds have been chosen appropriately, the inequality is very strict given the knowledge. To calculate the output intervals 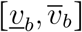 for 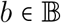, we can generate a modified circuit, which consists of 2*K* gates (if the circuit consists of *K*). This is done by generating two gates 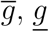 from one 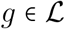 in the circuit, which contain the upper 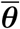 and lower 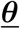 parametrizations. Then, for all following adjacent gates *g*′, we wire the output 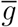 into 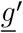 and 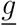 into 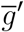. This resulting circuit then propagates input intervals 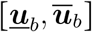 to output intervals 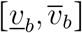. Once the output interval is calculated by standard signal propagation (see 4.4.1) through the modified circuit, the score (10) can be approximated by (11), taking the smallest difference 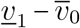. The generation of 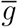 and 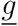 can thereby be done offline in advance and the new information can be gathered in an extended gate library.

As a small addition, and to give an idea of possible further considerations, we also propose a relaxed, less strict version of this score. Since it is easy to calculate output interval bounds 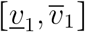 associated with Boolean 1 and 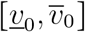 associated with Boolean 0, we can again think of these intervals as supporting output distributions. We could e.g. use this as a starting point for approximations of (8). Doing so, a reasonable assumption – if nothing else than the interval boundaries were known – would be assuming maximum entropy and therefore two uniform distributions with support within the interval boundaries. These can then again be scored using e.g. the E-Score (8).

The source code of the proposed synthesis and scoring methods is available at https://www.rs.tu-darmstadt.de/ARCTIC.

## 4.6 Supporting Information

(A) Pseudo code algorithms of the enumeration of structural circuit variants and the generation of equivalent envelope-free circuits (B) Circuit diagrams of designs synthesized using structural variants and uncertainty-aware assignment optimization

## Supporting Information

### A Algorithms

#### A.1 Enumeration of Structural Circuit Variants

The following pseudo codes depict the enumeration and pruning procedure for synthesizing structural circuit variants and its recursive enumeration kernel.

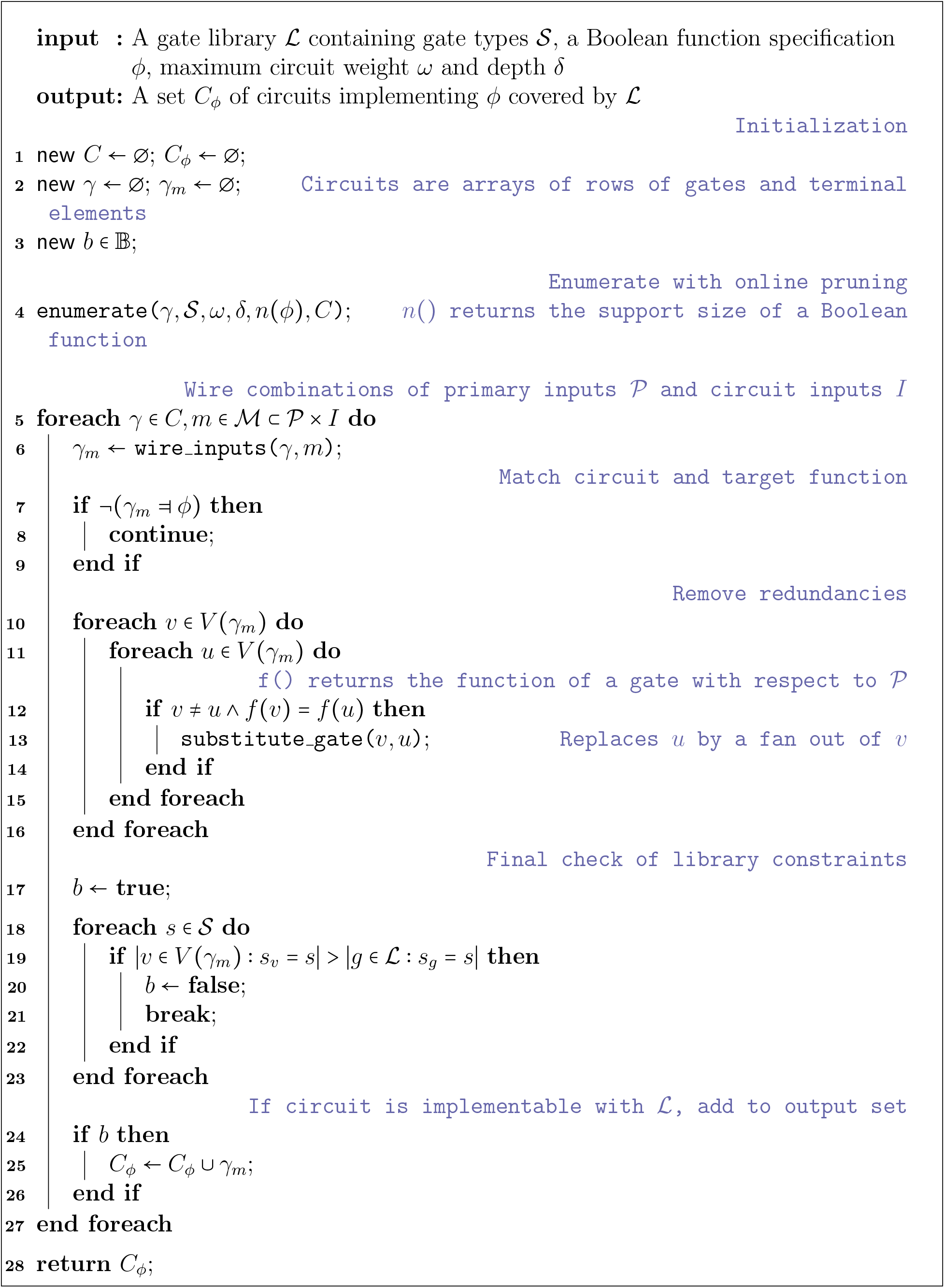

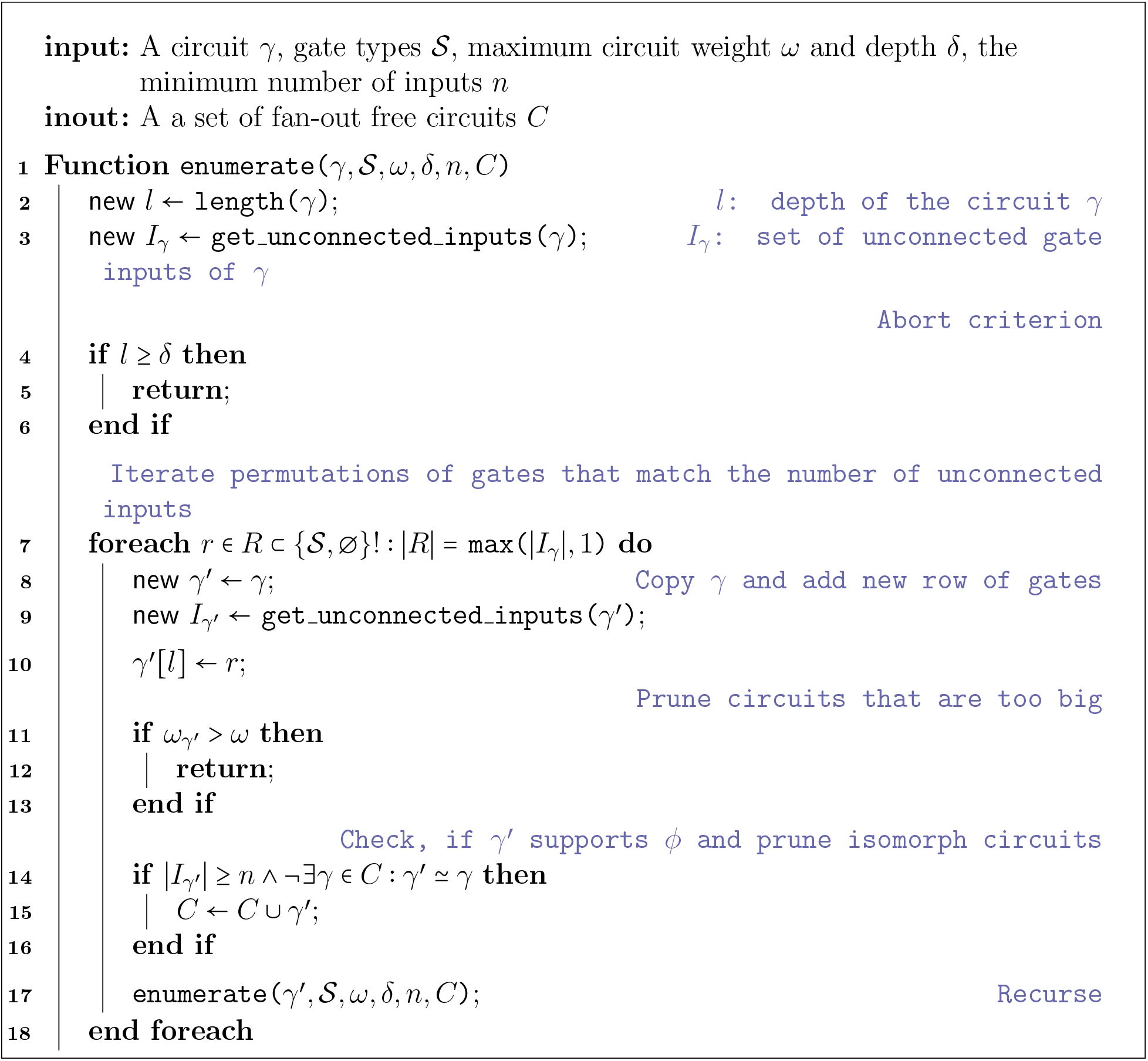

### A.2 Generation of an Equivalent Envelope-Free Circuit

The equivalent envelope-free circuit is just a ‘common’ circuit *C*^∗^, which is capable of carrying out the propagation of intervals through an original circuit *C*. Exploiting the monotonicity of all gate transfer functions in an extended gate library 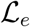, which contains tuples 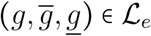 for each 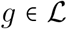, the circuit *C*^∗^ contains twice as many gates, only twice as many edges and its result is valid on the whole input domain.

For details on envelopes and the interval-based scoring, please refer to the Methods section from the original manuscript.

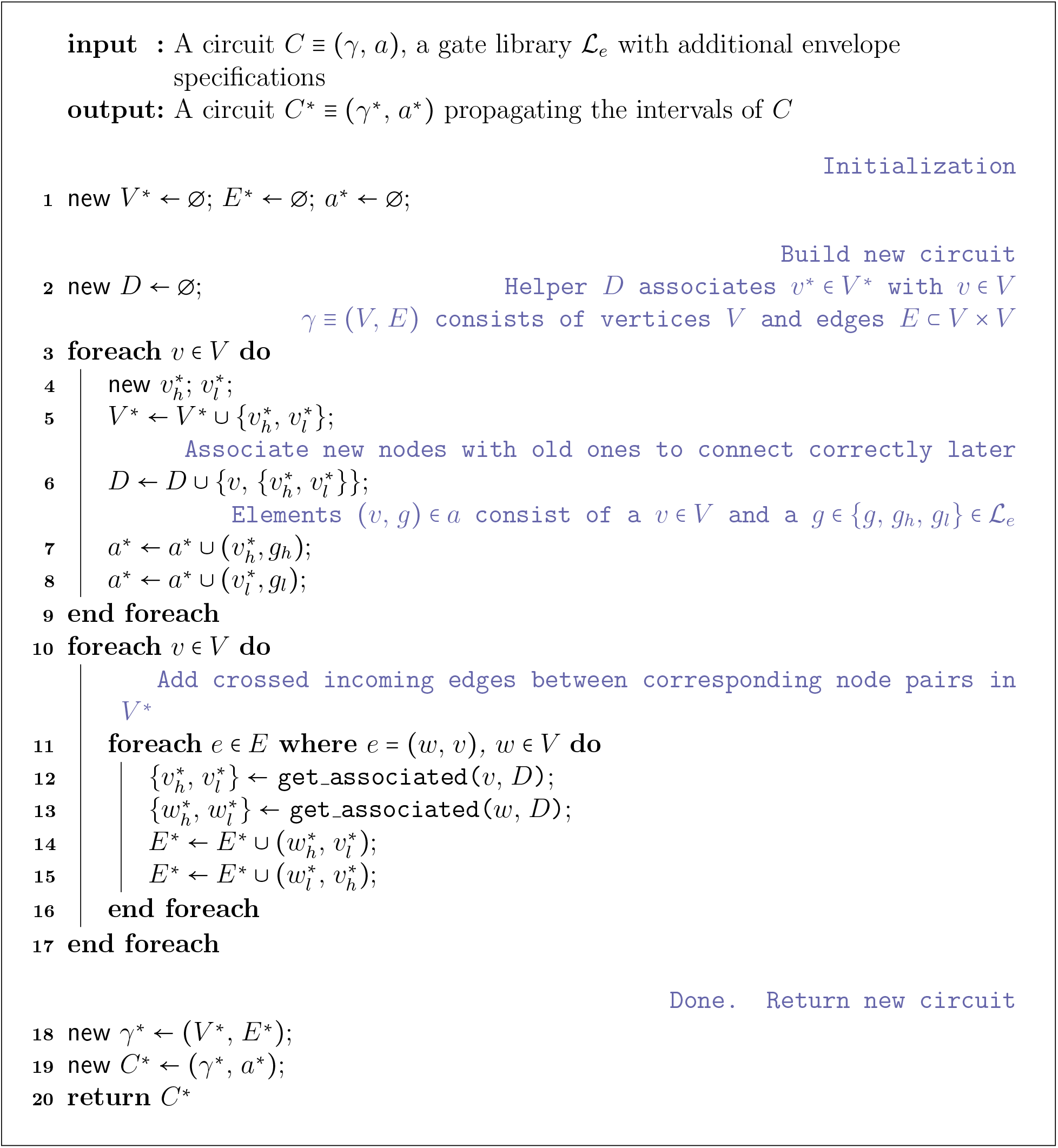

### B Synthesized Circuit Designs

#### B.1 Structural Variants, Classical Assignment Optimization

In the following, circuits synthesized by Cello and their structural variants synthesized by the proposed method are depicted, together with the optimal gate assignment found using the Cello score. Their corresponding final Cello scores are written below each. The diagrams have been automatically generated from the synthesis results.

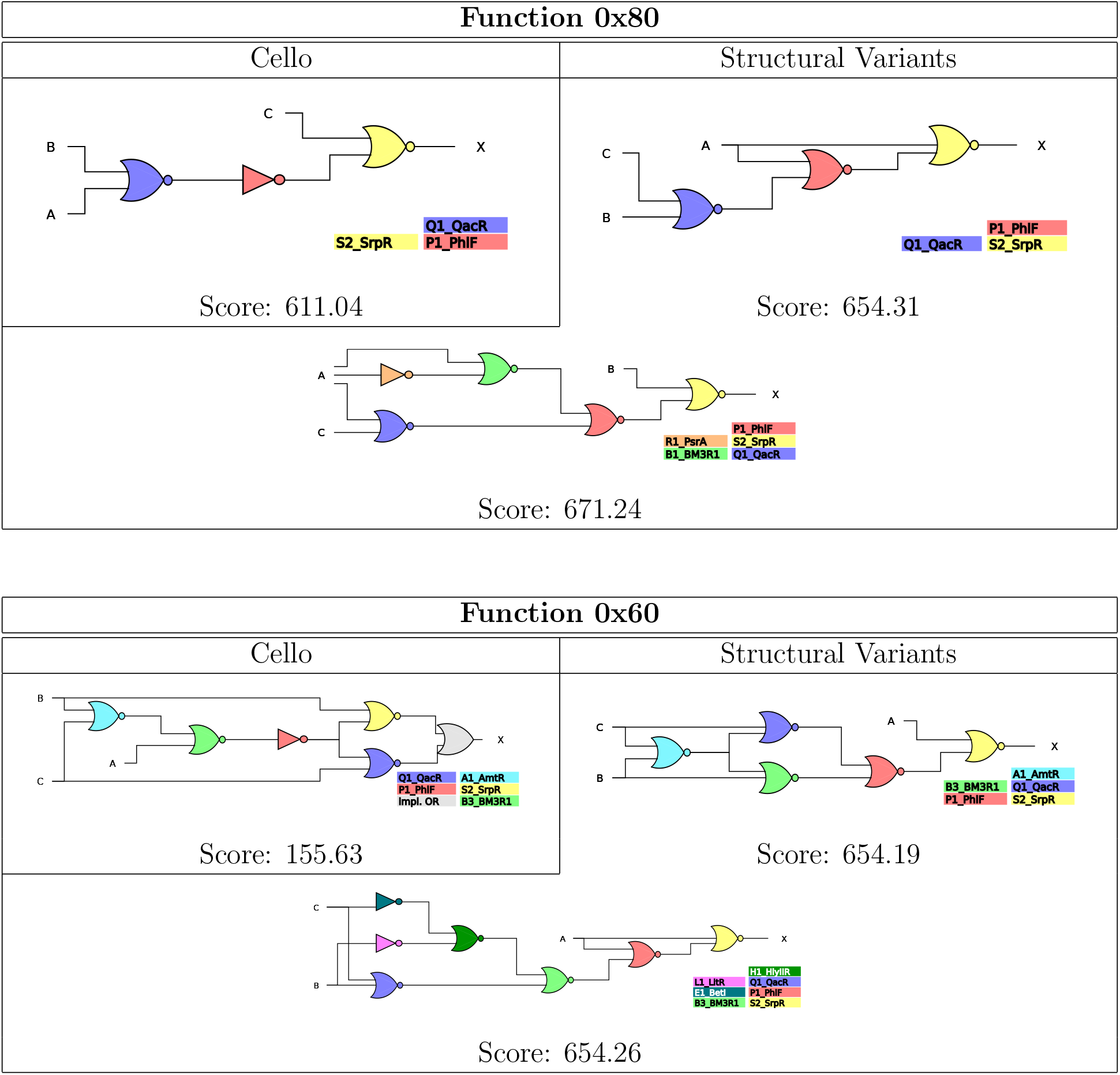

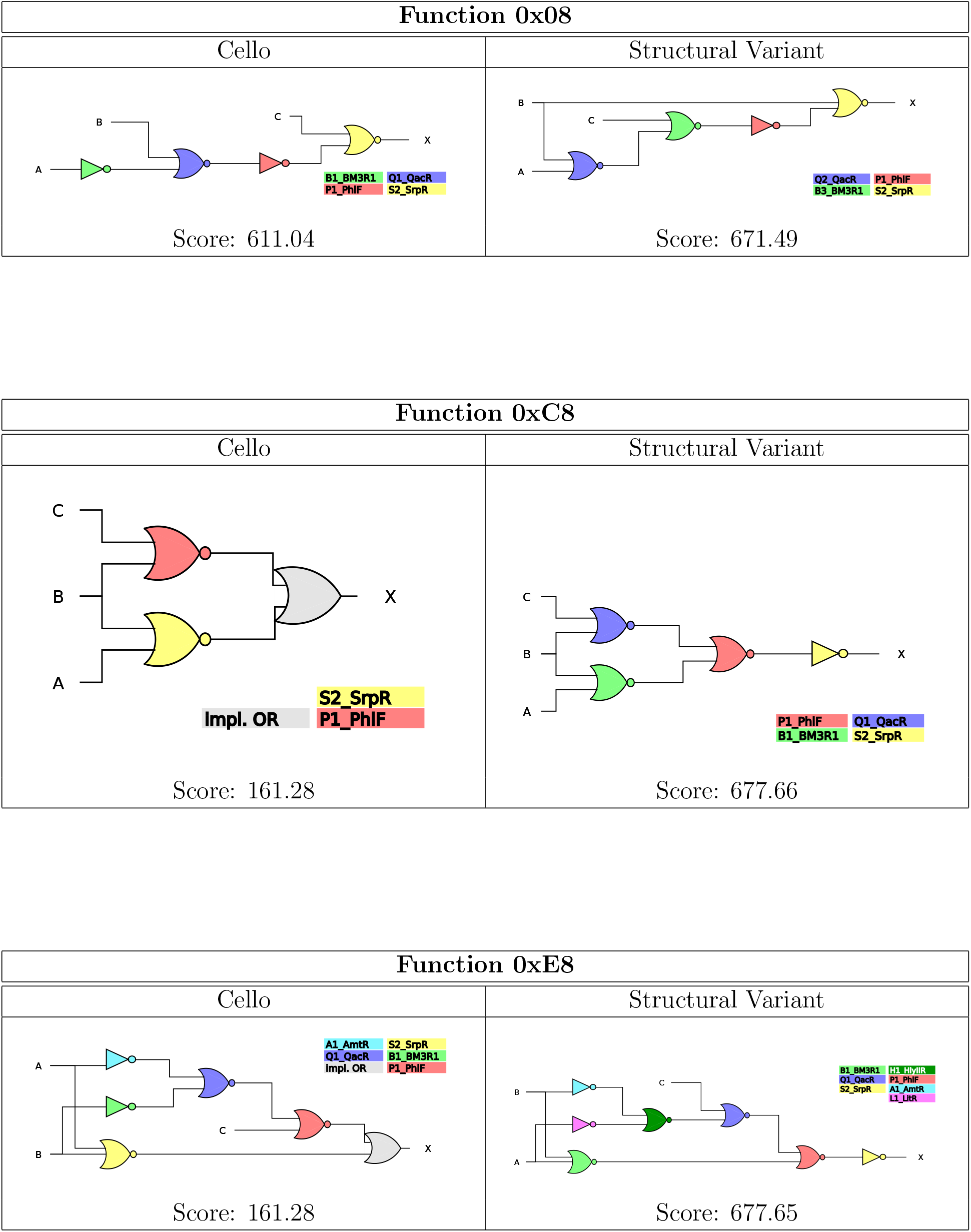

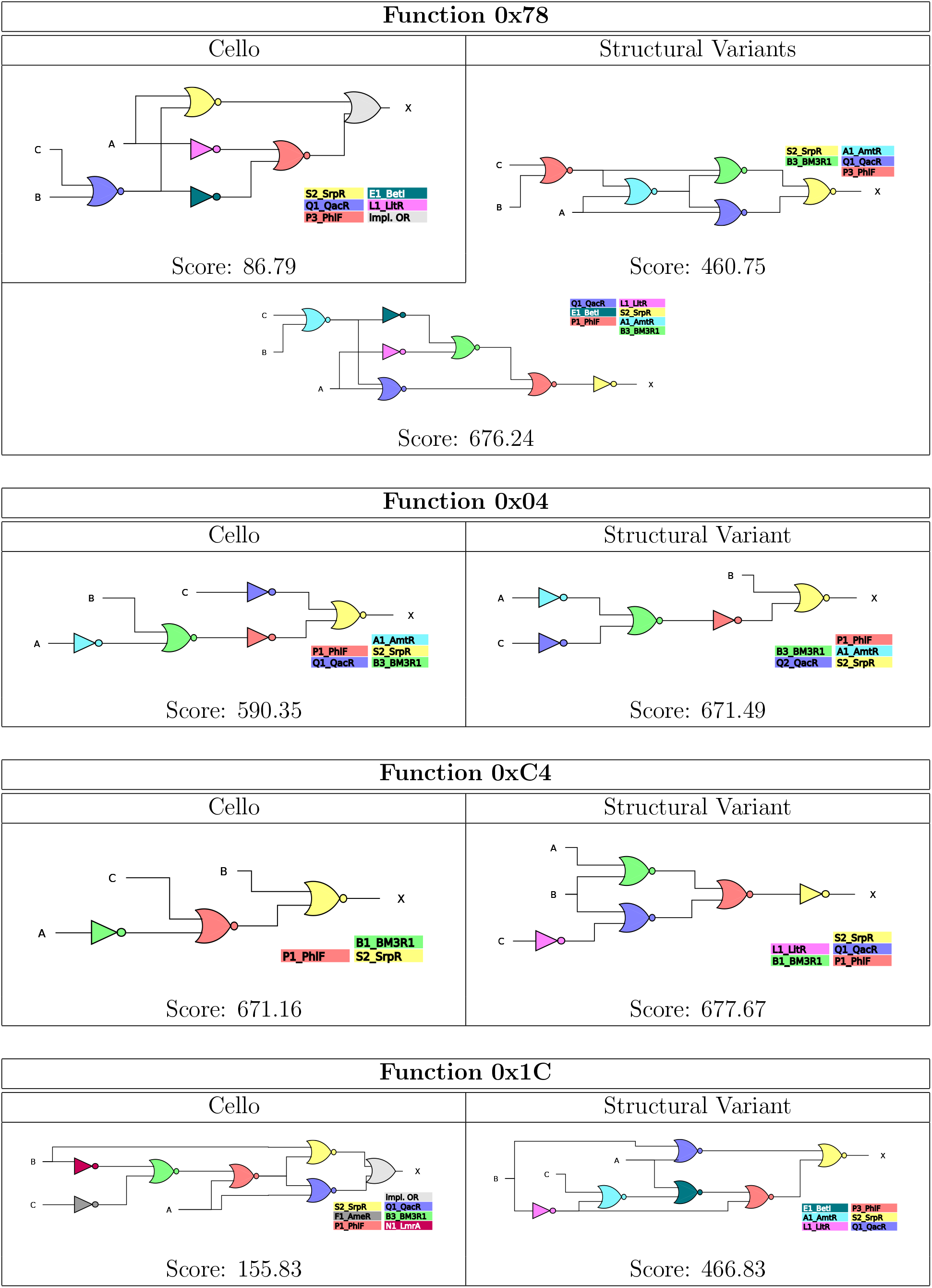

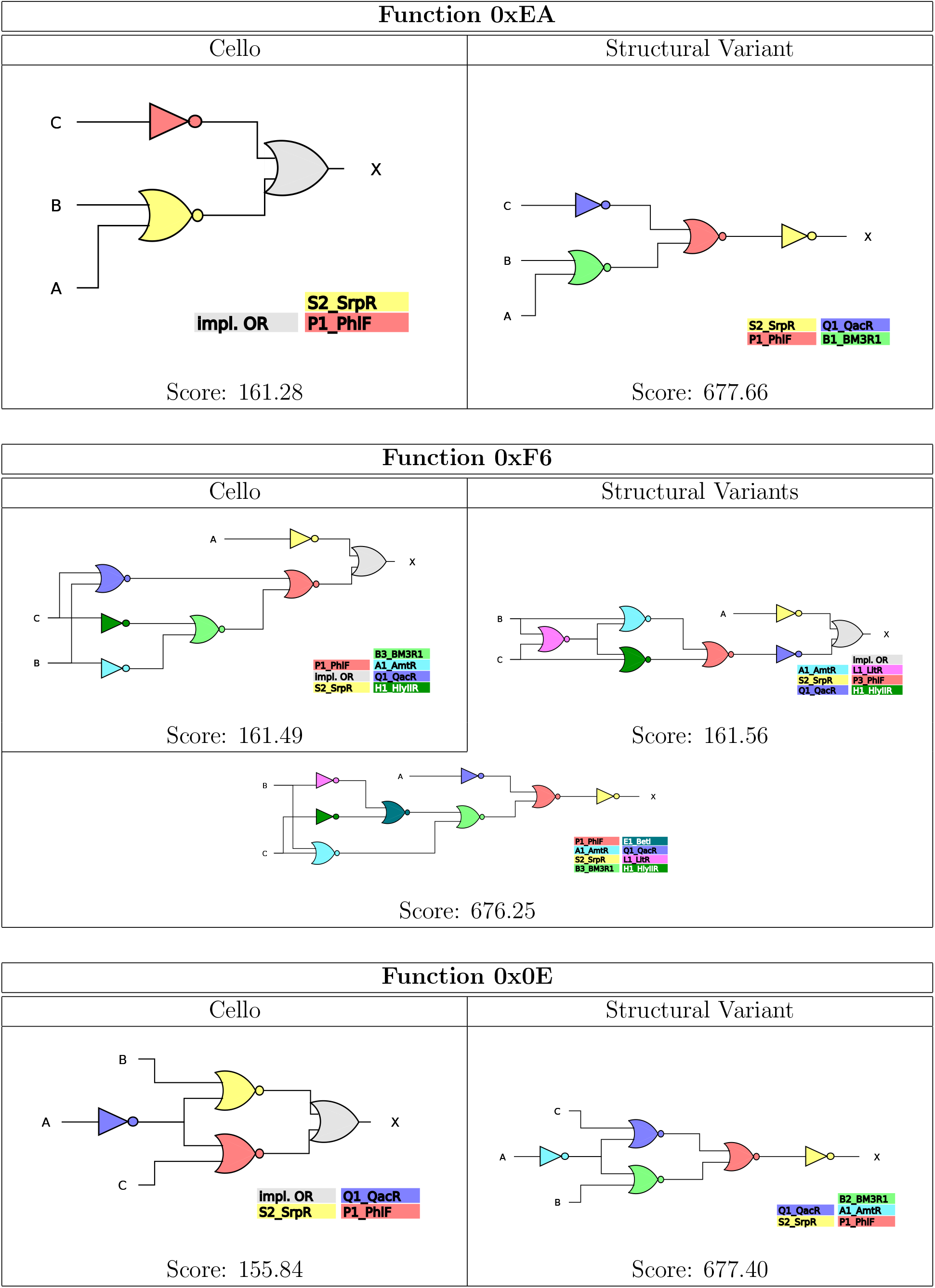

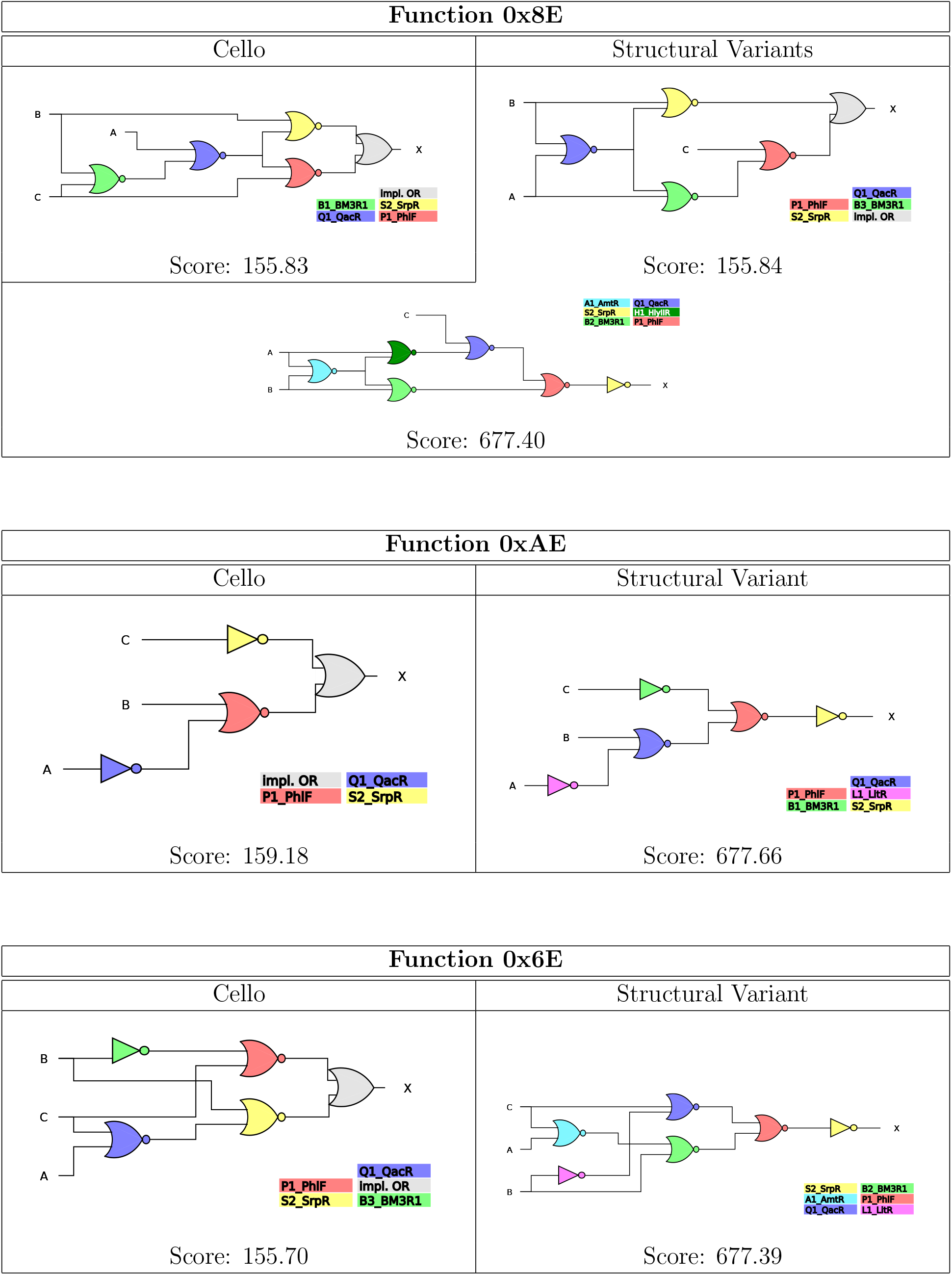

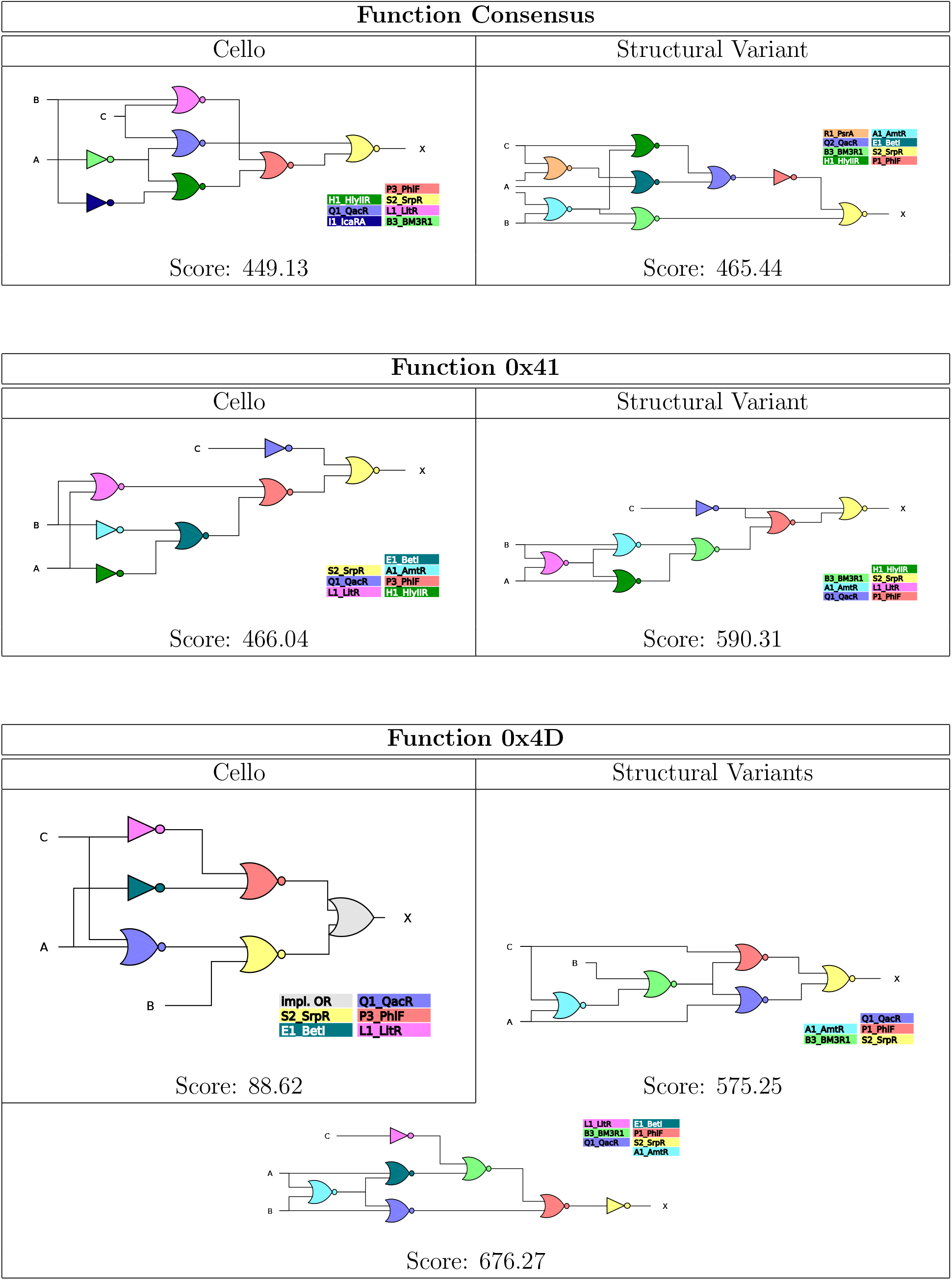

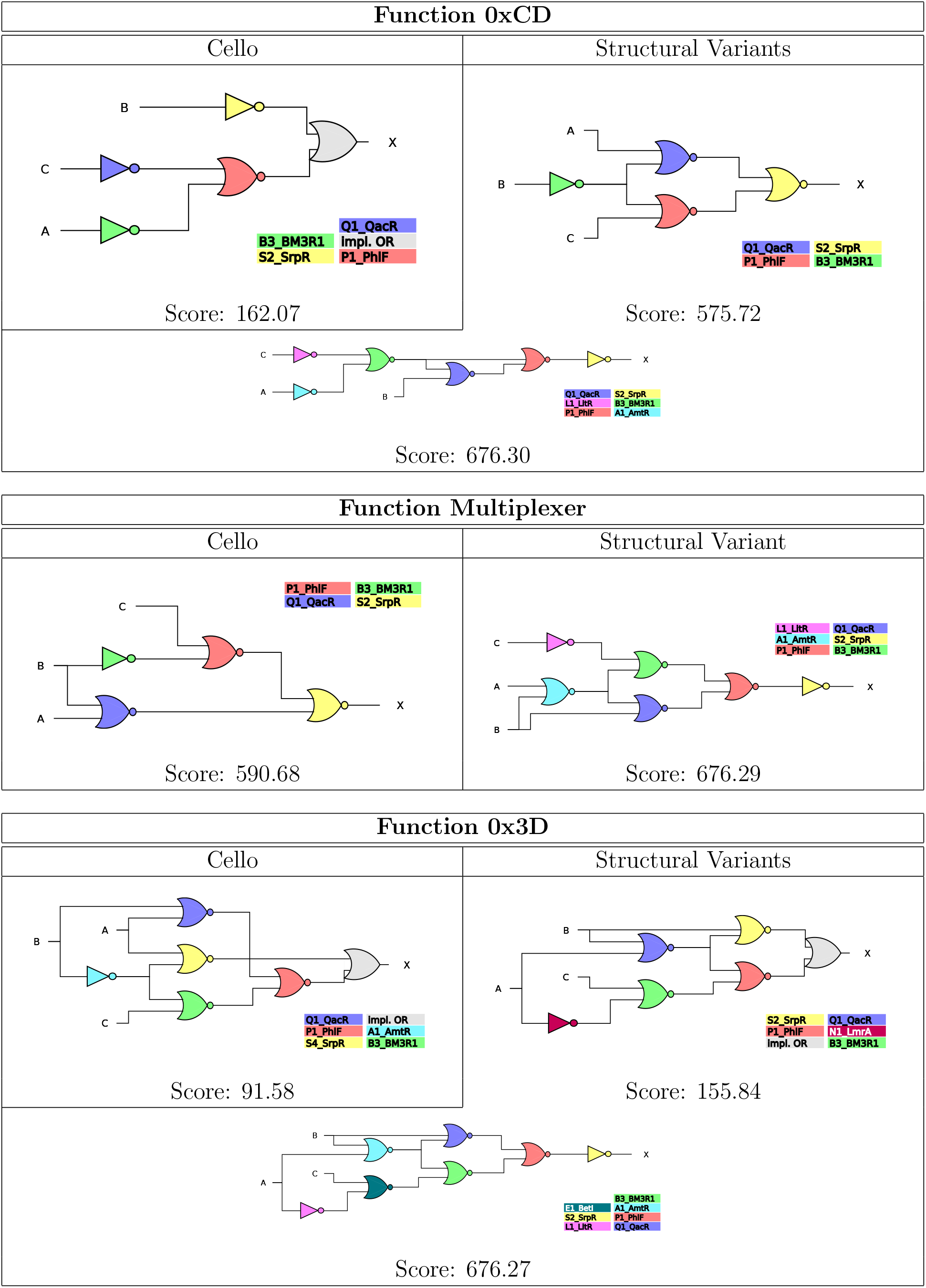

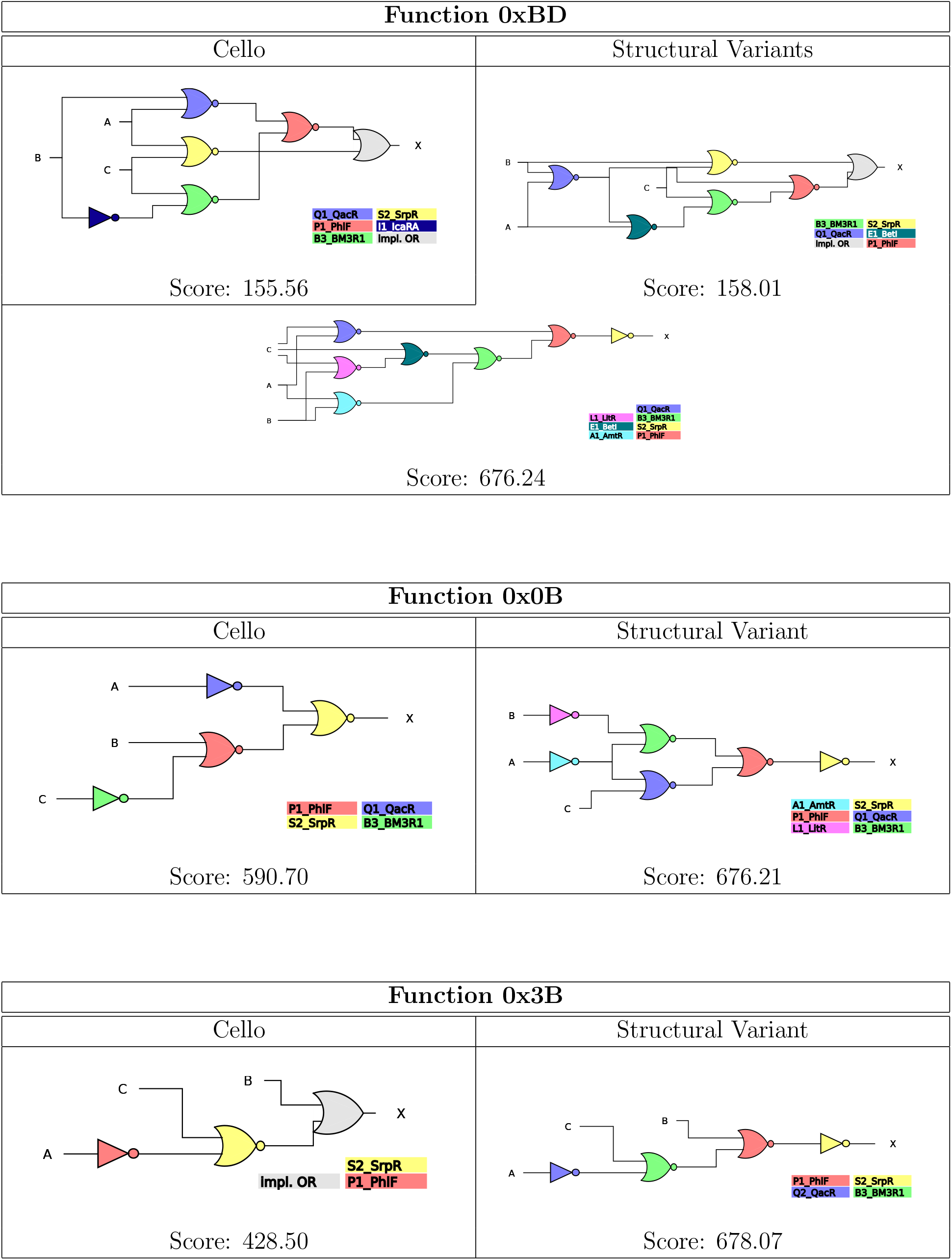

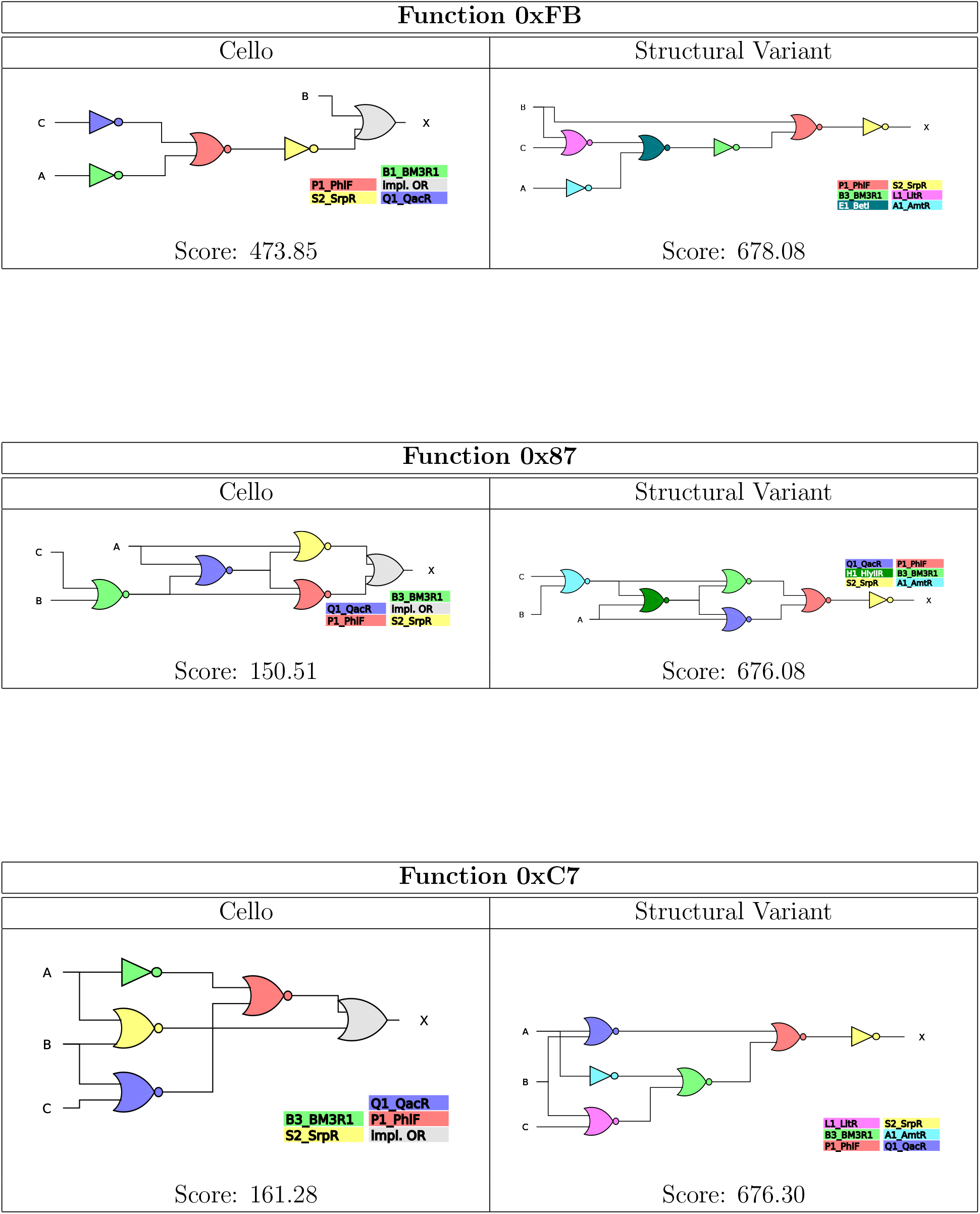

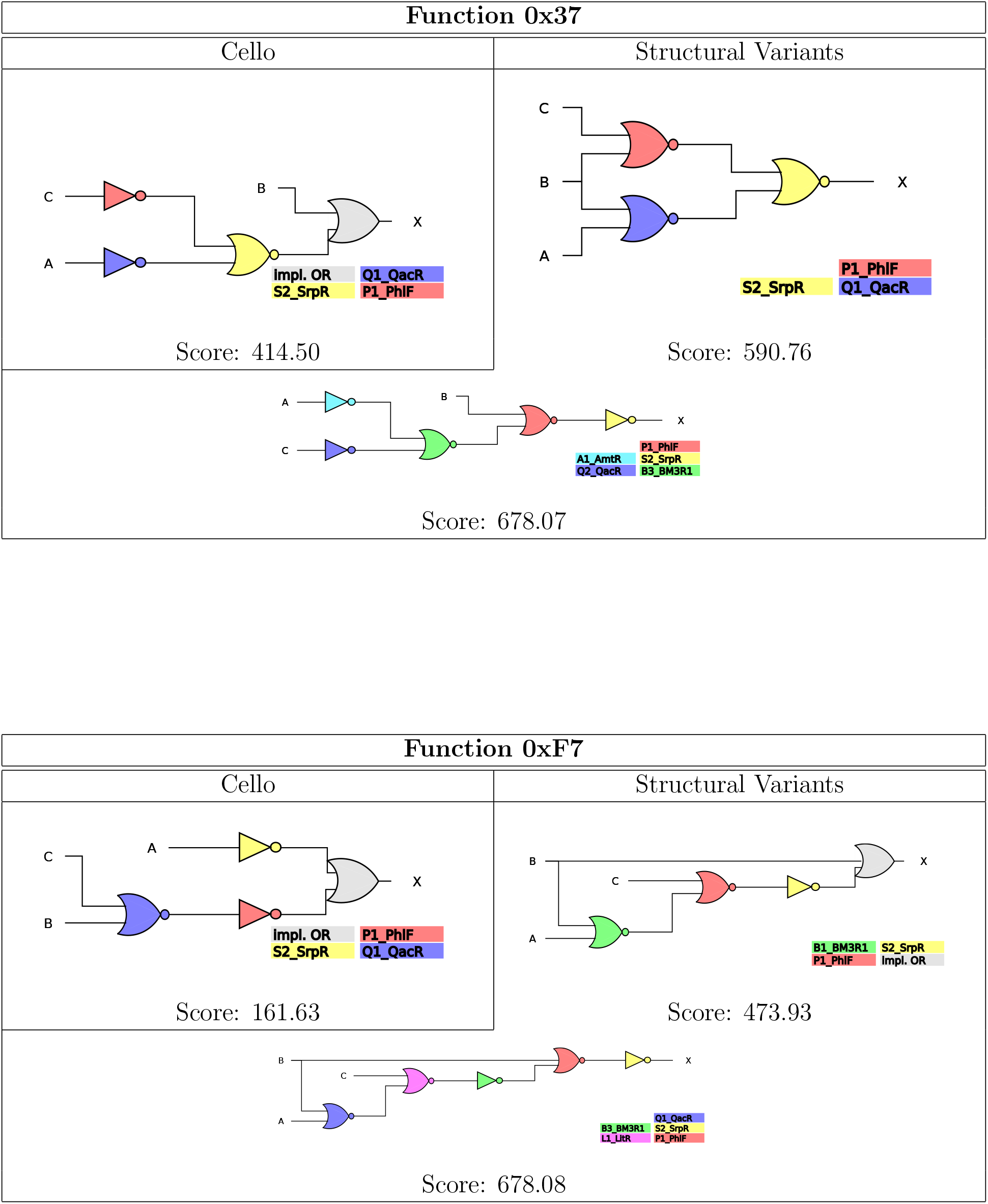

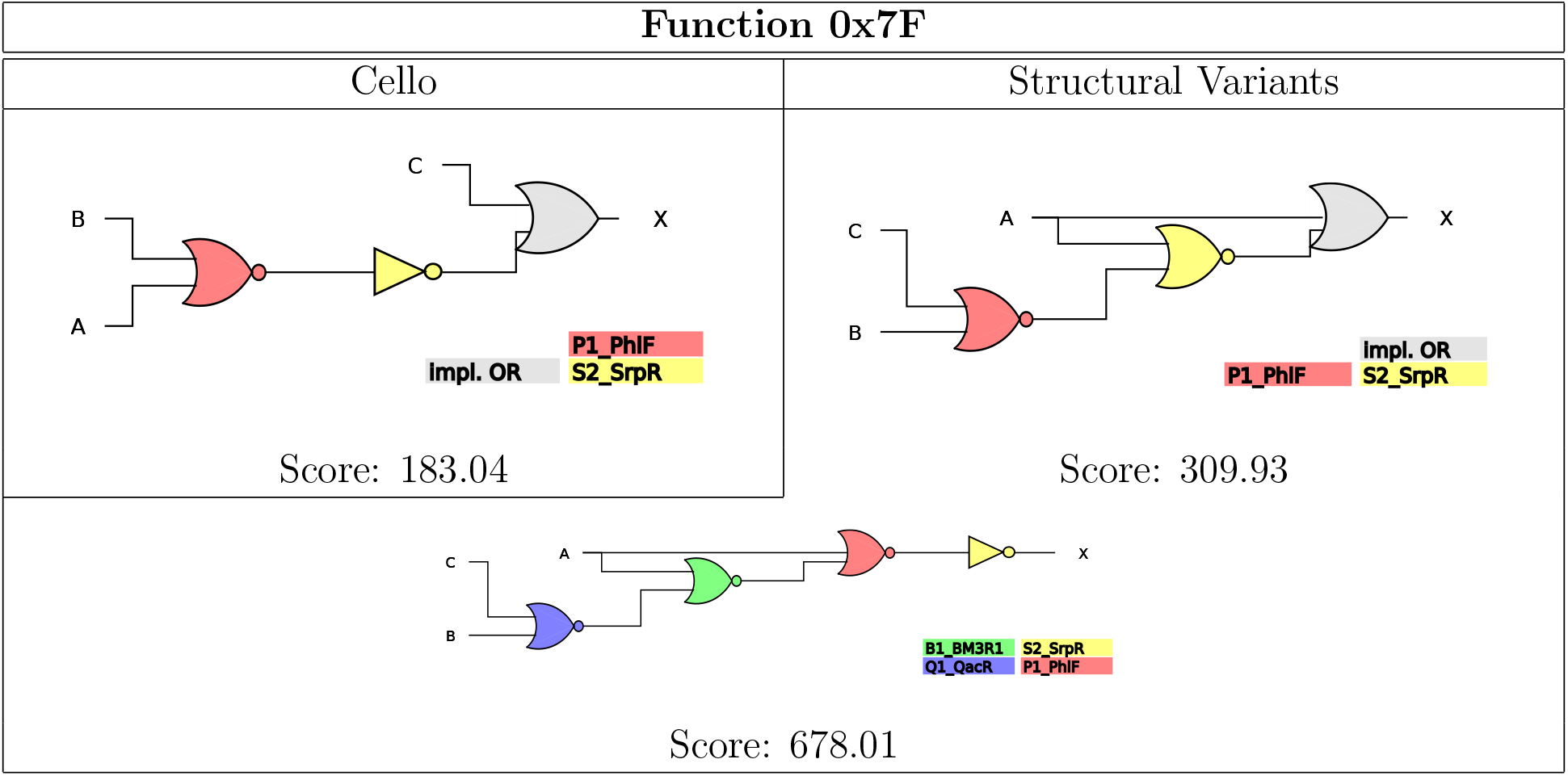

#### B.2 Classical Structure, Uncertainty-Aware Assignment Optimization

In the following, the three circuits mentioned in the main text 0x1c, 0x81 and 0x41 synthesized by Cello (so the non-modified original circuit structure) are depicted together with the optimal gate assignment found using the Cello score and the expectation-based score. The least separated on and off output histograms and their resulting final Cello and expectation-based scores are written below each.

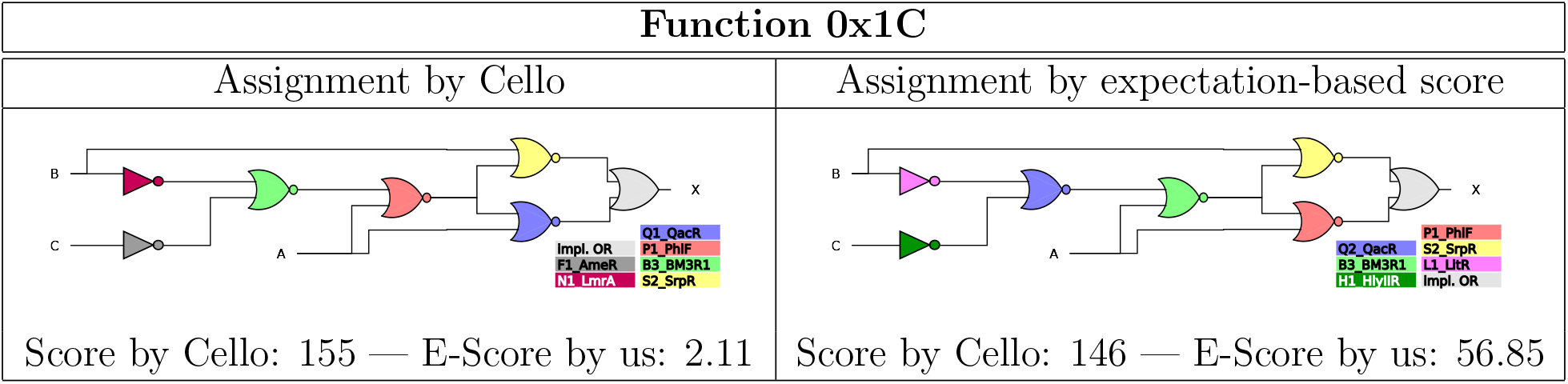

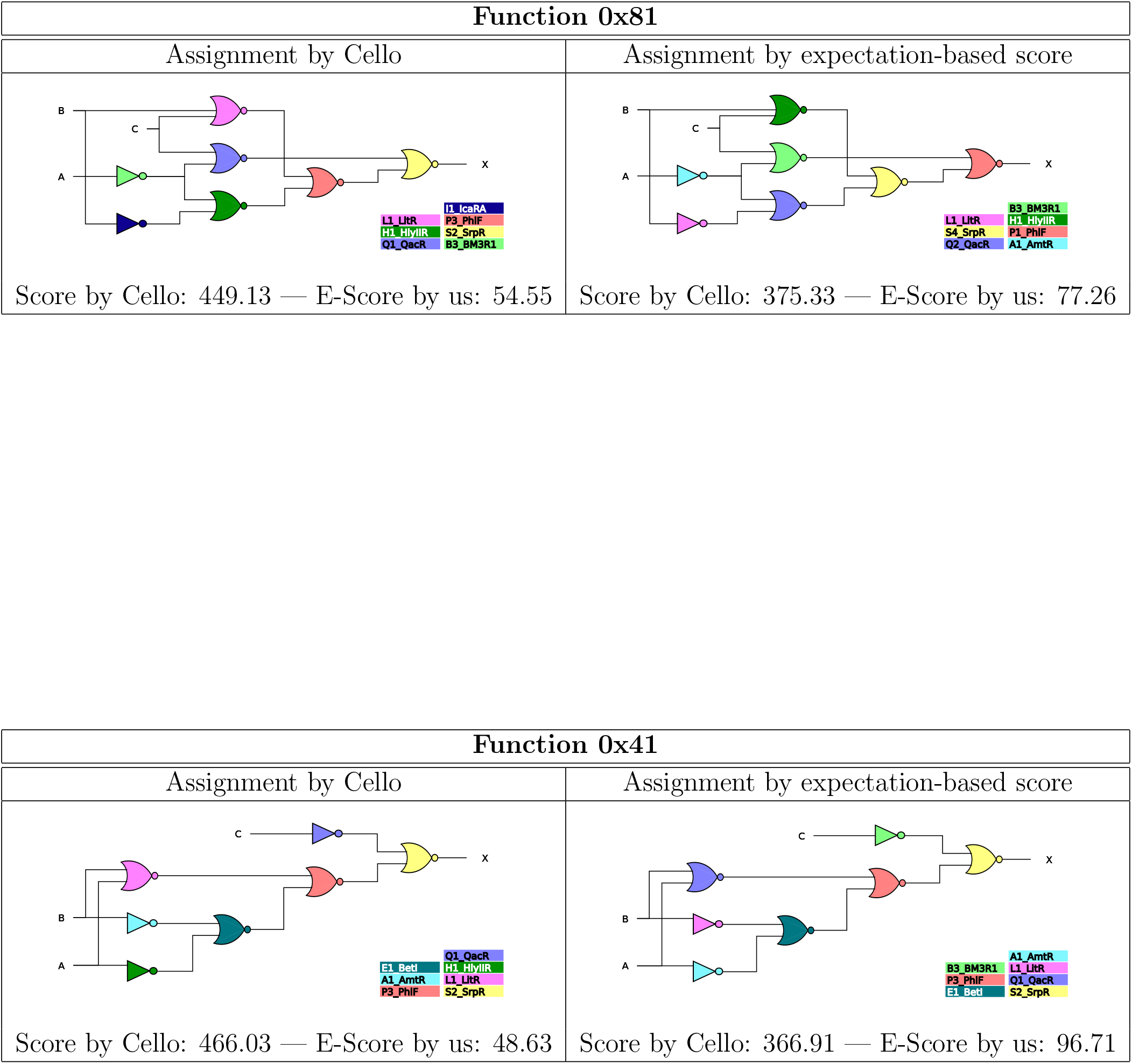

